# Transcriptome Dynamics of Floral Organs Approaching Blooming in the Flowering Cherry (*Cerasus* × *yedoensis*) Cultivar ‘Somei-Yoshino’

**DOI:** 10.1101/2021.10.26.465862

**Authors:** Kenta Shirasawa, Tomoya Esumi, Akihiro Itai, Sachiko Isobe

**Affiliations:** Laboratory of Plant Genetics and Genomics, Department of Frontier Research and Development, Kazusa DNA Research Institute, Kisarazu, Chiba 292-0818, Japan; Laboratory of Pomology & Viticulture, Department of Agricultural and Life Sciences, Shimane University, Matsue, Shimane 690-8504, Japan; Laboratory of Plant Resource Science, Department of Agricultural and Life Science, Kyoto Prefectural University, Sakyo, Kyoto 606-8522, Japan

**Keywords:** cherry blossom forecast, flowering cherry, RNA sequencing, time-course analysis, transcriptomics

## Abstract

To gain insights into the genetic mechanisms underlying blooming and petal movement in flowering cherry (*Cerasus* × *yedoensis*), we performed time-course RNA-seq analysis of the floral buds and open-flowers of the most popular flowering cherry cultivar, ‘Somei-Yoshino’. Independent biological duplicate samples of floral buds and open-flowers were collected from ‘Somei-Yoshino’ trees grown at three different locations in Japan. RNA-seq reads obtained from floral bud and open-flower samples collected in the current study (in 2019) and in a previous study (in 2017) were aligned against the genome sequence of ‘Somei-Yoshino’ to quantify gene transcript levels. Clustering analysis of RNA-seq reads revealed dynamic changes in the transcriptome, with genes in seven modules predominantly expressed at specific time points, ranging from 5 weeks before flowering to 2 weeks after flowering. Based on the identified gene modules and Gene Ontology (GO) terms enriched at different floral stages, we speculate that the genetic mechanisms underlying petal movement and flower opening in cherry involve the processes of development, cell wall organization, reproduction, and metabolism, which are executed by genes encoding transcription factors, phytohormones, transporters, and polysaccharide metabolic enzymes. Furthermore, we propose a method for cherry bloom forecasting, based on gene expression levels at different time points before flowering as RNA markers.

## 1 Introduction

Flowering cherry, also known as sakura, typically blooms in the spring and is valued as a popular ornamental flower across the world. ‘Somei-Yoshino’ (*Cerasus* × *yedoensis*), which is presumed to be an interspecific hybrid between *C. spachiana* and *C. speciosa* (Takenaka, 1963; Innan et al., 1995; Nakamura et al., 2015), is the most popular cultivar of flowering cherry in Japan. Given its genomic heterozygosity and self-incompatibility, ‘Somei-Yoshino’ is propagated by grafting (Iketani et al., 2007). Because of its clonal nature, ‘Somei-Yoshino’ trees planted within a specific location bloom at the same time; however, across the Japanese archipelago, the blooming front progresses from south to north because of differences in environmental conditions. Since the blooming date is important for the tourism industry in the spring season, forecasting methods based on cumulative temperature have been developed to predict the flowering date of cherry blossoms (Aono and Murakami, 2017).

Flowering involves two main processes: floral bud initiation and flower opening. The molecular mechanisms of floral bud initiation have been well studied in *Arabidopsis thaliana* and rice (*Oryza sativa*) as flowering models of long- and short-day plants, respectively, revealing that *FLOWERING LOCUS T* (*FT*) is the key gene involved in floral bud differentiation (Izawa et al., 2003). This mechanism is widely conserved across plant species (Higuchi, 2018). In woody plants, following floral bud initiation, buds enter a period of dormancy, which is followed by dormancy release and bud break. In the Rosaceae family members, *DORMANCY-ASSOCIATED MADS-box* (*DAM*) genes have been reported to regulate bud dormancy (Yamane, 2014). However, while the physiological aspect of the flower opening mechanism has been thoroughly investigated, only a few studies have been conducted to explore the genetic basis of this mechanism (van Doorn and Van Meeteren, 2003).

The genome sequence of ‘Somei-Yoshino’ has been determined at the chromosome level, and genes involved in the regulation of dormancy and flowering time have been identified in this cultivar through time-course transcriptome analysis (Shirasawa et al., 2019). Since expression levels of key genes transmit the environmental conditions, such as day-length and temperature, to biological processes such as blooming, the identification of these genes might help predict the blooming date in flowering cherry. Because the transcriptome is affected by environmental conditions (e.g., changes in weather and habitat), biological replications of flowering cherry trees over multiple years and locations would be required for the accurate identification of genes affecting the blooming date.

In this study, we aimed to identify genes uniquely expressed before and after flowering, and to obtain insights into the molecular mechanisms underlying blooming in flowering cherry. We collected floral buds and open-flowers from ‘Somei-Yoshino’ trees planted at three locations in Japan (Chiba, Kyoto, and Shimane) in 2019, and analyzed their transcriptome using the RNA sequencing (RNA-seq) technology. The time-course transcriptome data generated in the current study and in our previous study (Shirasawa et al., 2019) was used to characterize gene expression patterns in ‘Somei-Yoshino’ floral buds, some of which could be used to forecast the blooming time and to gain insights into the molecular mechanisms controlling blooming in flowering cherry.

## 2 Materials and Methods

### 2.1 Plant Materials

Seven clonal trees of ‘Somei-Yoshino’ were used in this study. One tree was planted at the Kazusa DNA Research Institute (KDRI; Kisarazu, Chiba, Japan), and three trees each were planted at two different locations: Shimane University (SU; Matsue, Shimane, Japan; trees 1–3) and Kyoto Prefectural University (KPU; Sakyo, Kyoto, Japan; trees A–C). Floral buds and open-flowers were collected in 2019 over an extended time period, ranging from 36 days before flowering (DBF) to 11 days after flowering (DAF) (Figure 1), corresponding to 6 weeks before flowering (WBF) to 2 weeks after flowering (WAF) (Supplementary Table S1).

**Figure 1.**
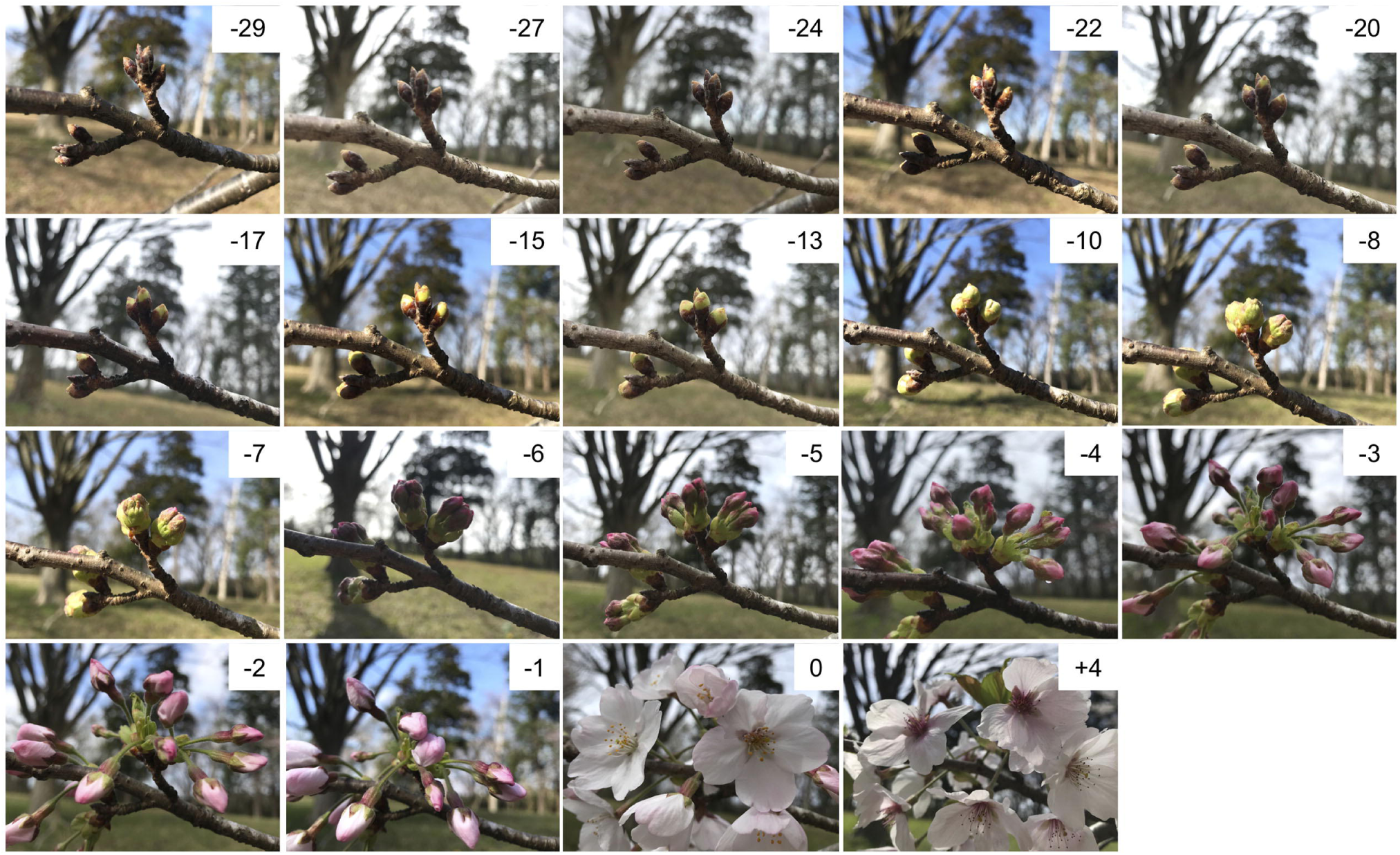
Floral buds and open-flowers of ‘Somei-Yoshino’. The numbers in each picture indicate days before (-) or after (+) the flowering day (0). Photos were taken at Kazusa DNA Research Institute in 2021.

### 2.2 RNA-seq Analysis

Library preparation and sequencing analysis were performed as described in Shirasawa et al. (2019). In short, total RNA was extracted from the buds and open-flowers with RNeasy Plant Min Kit (Qiagen, Hilden, Germany). The RNA was treated with RNase-free DNase (Promega, Madison, WI, USA) and used for library construction with TruSeq Stranded mRNA Library Prep Kit (Illumina, San Diego, CA, USA). The library was sequenced on NextSeq 500 (Illumina) in paired-end, 76 bp mode.

### 2.3 RNA Sequence Data Processing

RNA-seq data was analyzed as described previously (Shirasawa et al., 2019). High-quality reads were selected by trimming the adapter sequences using fastx_clipper (parameter, -a AGATCGGAAGAGC) in the FASTX-Toolkit v.0.0.13 (http://hannonlab.cshl.edu/fastx_toolkit), and by deleting low-quality bases using PRINSEQ v.0.20.4 (Schmieder and Edwards, 2011). High-quality reads were mapped to the CYE_r3.1_pseudomolecule sequence (Shirasawa et al., 2019) using HISAT2 v.2.1.0 (Kim et al., 2015), and reads mapped to each gene model were quantified and normalized to determine the number of fragments per kilobase of transcript per million mapped reads (FPKM) using StringTie v.1.3.5 (Pertea et al., 2015) and Ballgown v.2.14.1 (Frazee et al., 2015), as described previously (Pertea et al., 2016). Genes showing a variance of ≥1 in expression levels among samples were used for further analysis.

### 2.4 Gene Ontology (GO) Enrichment Analysis of Gene Modules

The weighted gene correlation network analysis (WGCNA; v.1.66) package of R (Langfelder and Horvath, 2008) was used for module detection. The floral bud and open-flower samples were grouped in accordance with gene expression levels using the hierarchical clustering algorithm. Then, highly co-expressed gene clusters were constructed as modules. Genes included in the modules were functionally annotated by performing a BLAST sequence similarity search (Altschul et al., 1997) against the non-redundant (nr) protein database. GO terms were assigned to genes, and the statistical analysis of the GO term enrichment in each module was performed using Fisher’s exact test implemented in OmicsBox (BioBam, Valencia, Spain). GO maps were drawn with QuickGO (Huntley et al., 2009).

## 3 Results

### 3.1 RNA-seq Analysis and Quantification of Gene Expression Levels

A total of 94 samples were collected from flowering cherry trees at KDRI, SU, and KPU over a period of 47 days (i.e., 36 days before anthesis to 11 days post-anthesis) in 2019 (Figure 1, Supplementary Table S1). Total RNA was extracted from the samples and subjected to RNA-seq analysis, which returned approximately 4.7 million reads per sample. Nucleotide sequence data of the RNA-seq were deposited in the DDBJ Sequence Read Archive (accession numbers DRA012801, DRA012802, and DRA012803). We also utilized the RNA-seq data generated previously (Shirasawa et al. 2019; DRA accession number DRA008100) from samples collected at KDRI in 2017. The RNA-seq reads from both datasets were mapped on to the genome sequence of ‘Somei-Yoshino’ (Shirasawa et al., 2019), and expression levels were normalized in all samples based on the total reads to obtain FPKM values for each gene across the samples. Of the 95,076 genes predicted in the ‘Somei-Yoshino’ genome (Shirasawa et al., 2019), a total of 29,712 genes (31.3%) were expressed across the different samples, with a variance of ≥1.

### 3.2 Gene Module Detection

On the basis of the expression patterns of 29,712 genes, one floral bud sample collected at 27 days before anthesis from the KDRI location was identified as an outlier and therefore was excluded from further analyses (Supplementary Figure S1). The remaining 93 samples were roughly clustered into 53 highly co-expressed gene modules, based on the expression patterns of 29,712 genes (Supplementary Figure S2), which were further classed into four groups, based on eigengenes (Supplementary Figure S3). Genes in 7 out of 53 modules were prominently expressed at specific days before or after anthesis (Figure 2): dark-red module (110 genes expressed at 4–5 weeks before flowering [WBF]), tan module (267 genes, 4 WBF), pink module (332 genes, 3 WBF), royal blue module (127 genes, 2–3 WBF), midnight blue module (176 genes, 1–2 WBF), black module (457 genes, at flowering days), and sky-blue module (72 genes, 1–2 weeks after flowering [WAF]).

**Figure 2.**
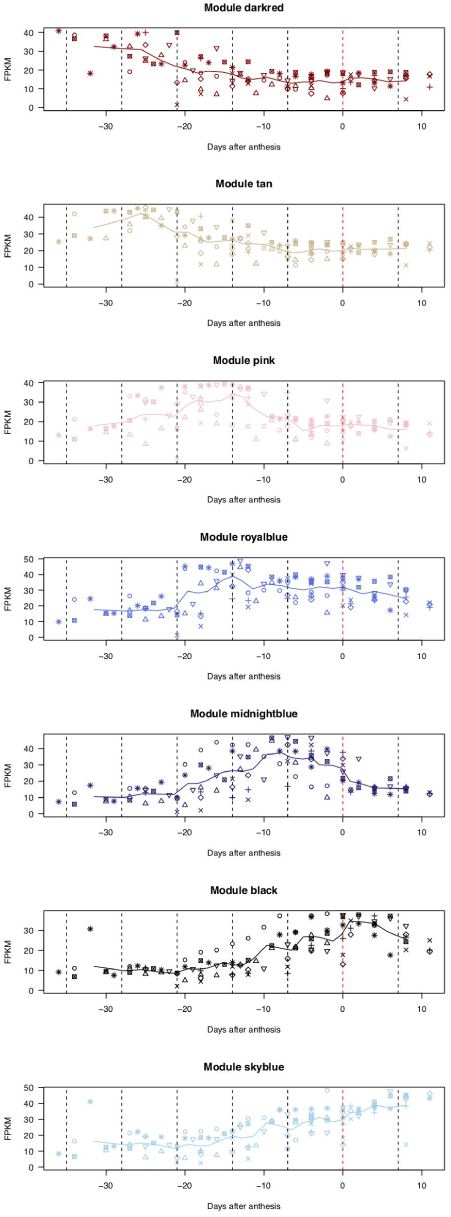
Expression patterns of genes categorized into seven modules. X-axis indicates the number of days before (-) or after (+) the flowering day (0). Lines show moving averages of gene expression (sliding window size = 8, walking speed = 4). Symbols indicate biological replicates: KDRI in 2017 (circle); KDRI in 2019 (triangle); SU1 in 2019 (plus); SU2 in 2019 (cross); SU3 in 2019 (diamond); KPUA in 2019 (upside-down triangle); KPUB in 2019 (square cross); and KPUC in 2019 (star).

### 3.3 GO Enrichment Analysis

To identify the GO terms enriched in each module, the ratios of GO terms in each gene module were compared with those of the remaining gene set (Table 1, Figure 2, Supplementary Table S2, Supplementary Figure S4).

**Table 1.**
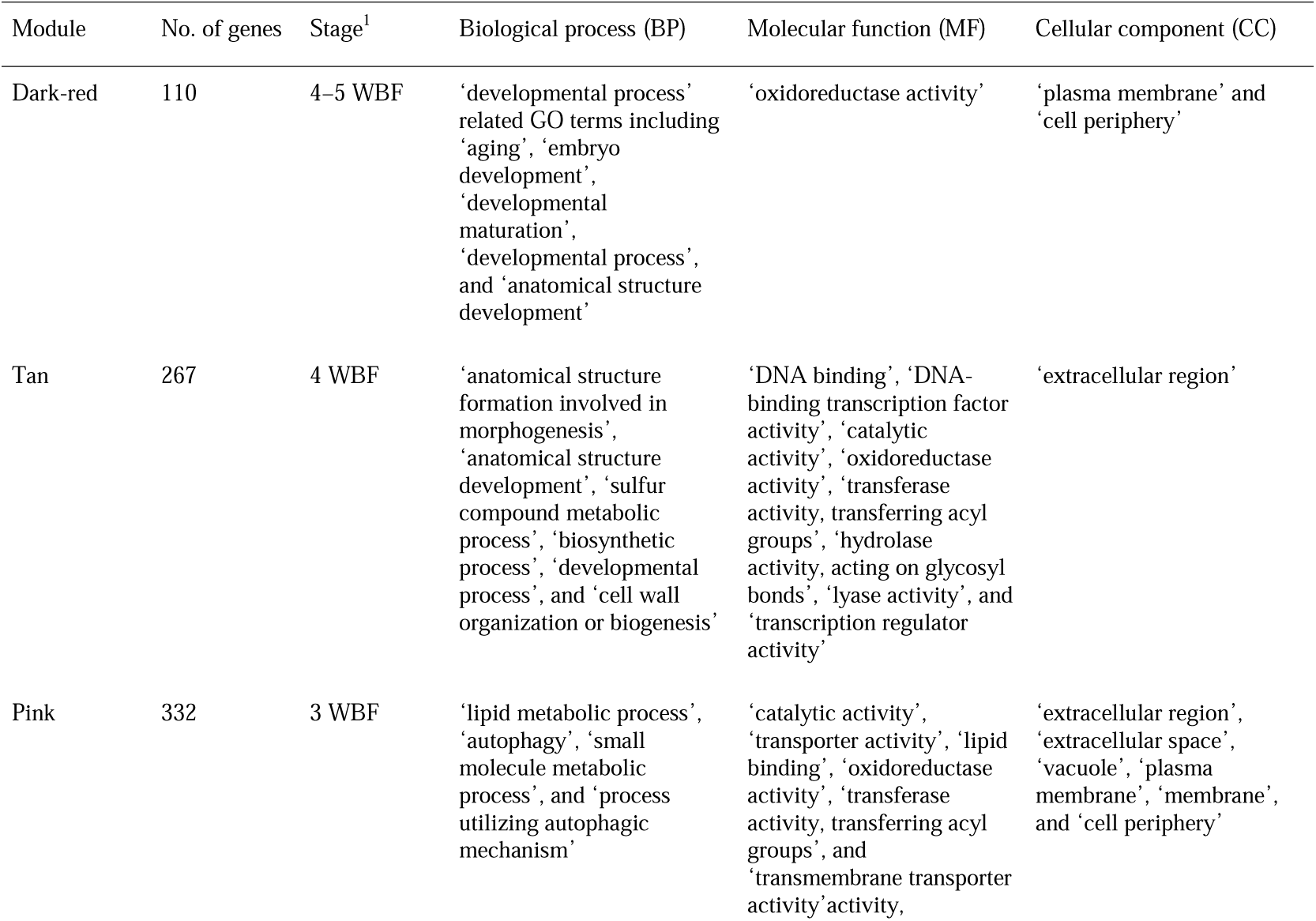
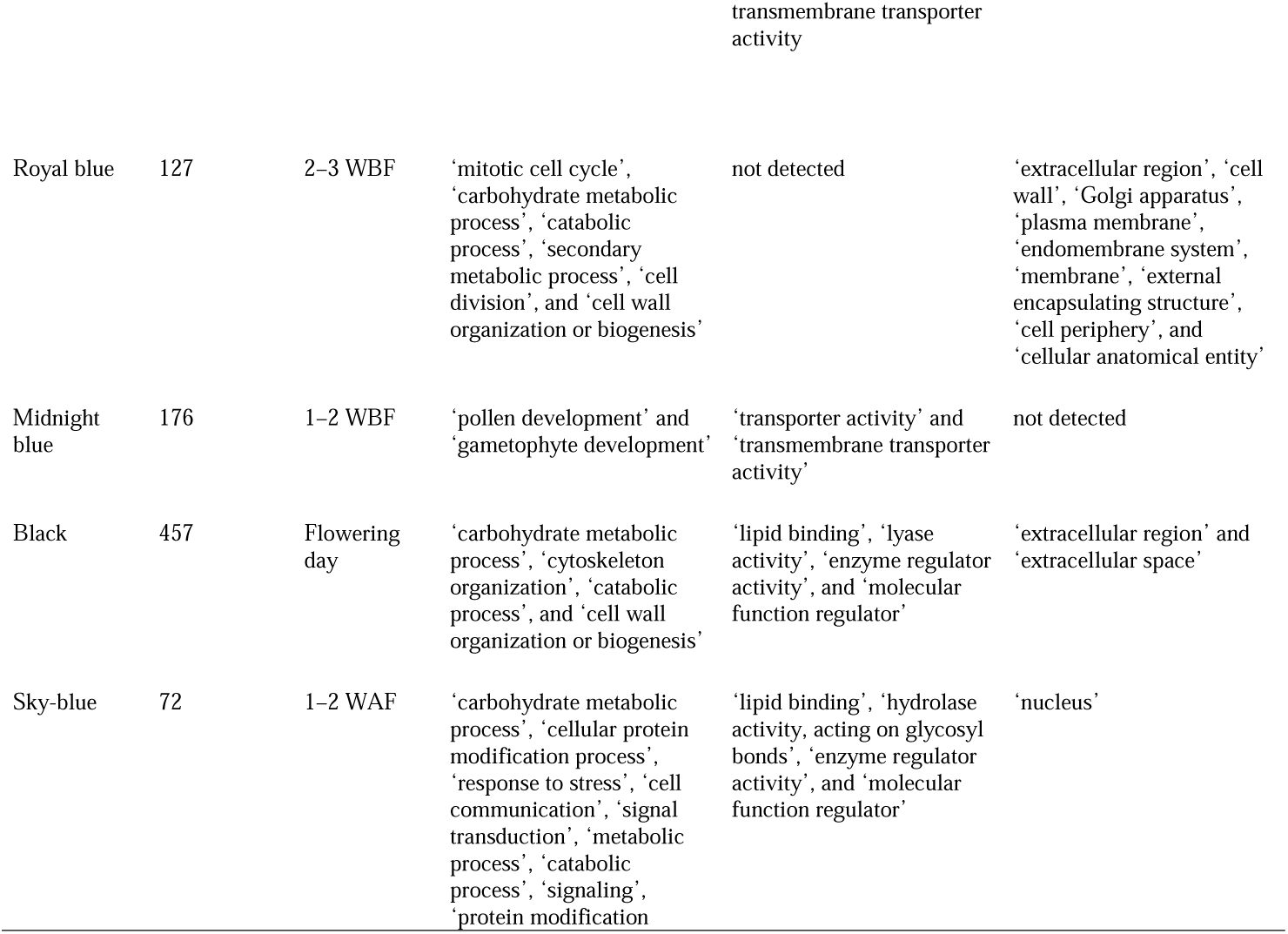
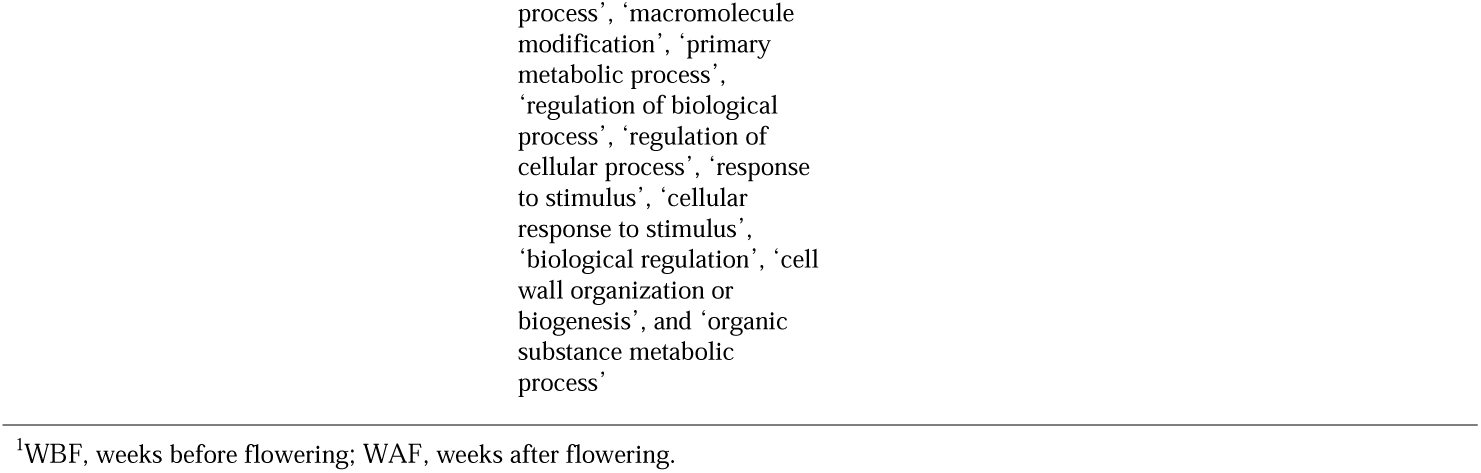
Gene Ontology (GO) terms enriched at the different flowering stages.

At 4–5 WBF (dark-red module), the following GO terms were enriched: biological process (BP) category: ‘developmental process’ related GO terms including ‘aging’, ‘embryo development’, ‘developmental maturation’, ‘developmental process’, and ‘anatomical structure development’; molecular function (MF) category: ‘oxidoreductase activity’; and cellular component (CC) category: ‘plasma membrane’ and ‘cell periphery’.

At 4 WBF (tan module), GO terms enriched in the BP category were not only related to ‘developmental process’ (‘anatomical structure formation involved in morphogenesis’ and ‘anatomical structure development’) but also ‘cellular process’ (‘sulfur compound metabolic process’, ‘biosynthetic process’, ‘developmental process’, and ‘cell wall organization or biogenesis’). GO terms enriched in the MF category included ‘DNA binding’, ‘DNA-binding transcription factor activity’, ‘catalytic activity’, ‘oxidoreductase activity’, ‘transferase activity, transferring acyl groups’, ‘hydrolase activity, acting on glycosyl bonds’, ‘lyase activity’, and ‘transcription regulator activity’, while those enriched in the CC category included ‘extracellular region’.

At 3 WBF (pink module), the following GO terms were enriched: BP category: ‘lipid metabolic process’, ‘autophagy’, ‘small molecule metabolic process’, and ‘process utilizing autophagic mechanism’; MF category: ‘catalytic activity’, ‘transporter activity’, ‘lipid binding’, ‘oxidoreductase activity’, ‘transferase activity, transferring acyl groups’, and ‘transmembrane transporter activity’; and CC category: ‘extracellular region’, ‘extracellular space’, ‘vacuole’, ‘plasma membrane’, ‘membrane’, and ‘cell periphery’.

At 2–3 WBF (royal blue module), the following GO terms were enriched: BP category: ‘mitotic cell cycle’, ‘carbohydrate metabolic process’, ‘catabolic process’, ‘secondary metabolic process’, ‘cell division’, and ‘cell wall organization or biogenesis’; and CC category: ‘extracellular region’, ‘cell wall’, ‘Golgi apparatus’, ‘plasma membrane’, ‘endomembrane system’, ‘membrane’, ‘external encapsulating structure’, ‘cell periphery’, and ‘cellular anatomical entity’.

At 1–2 WBF (midnight blue module), the enriched GO terms included ‘pollen development’ and ‘gametophyte development’ (BP category); and ‘transporter activity’ and ‘transmembrane transporter activity’ (MF category).

At the flowering stage (black module), the following GO terms were enriched: BP category: ‘carbohydrate metabolic process’, ‘cytoskeleton organization’, ‘catabolic process’, and ‘cell wall organization or biogenesis’; MF category: ‘lipid binding’, ‘lyase activity’, ‘enzyme regulator activity’, and ‘molecular function regulator’; and CC category: ‘extracellular region’ and ‘extracellular space’.

At 1–2 WAF (sky-blue module), the following GO terms were enriched: BP category: ‘carbohydrate metabolic process’, ‘cellular protein modification process’, ‘response to stress’, ‘cell communication’, ‘signal transduction’, ‘metabolic process’, ‘catabolic process’, ‘signaling’, ‘protein modification process’, ‘macromolecule modification’, ‘primary metabolic process’, ‘regulation of biological process’, ‘regulation of cellular process’, ‘response to stimulus’, ‘cellular response to stimulus’, ‘biological regulation’, ‘cell wall organization or biogenesis’, and ‘organic substance metabolic process’; MF category: ‘lipid binding’, ‘hydrolase activity, acting on glycosyl bonds’, ‘enzyme regulator activity’, and ‘molecular function regulator’; and CC category: ‘nucleus’. Thus, the GO terms enriched at this stage were drastically different from those enriched at the other stages.

### 3.4 Genetic Mechanisms Underlying Blooming in Flowering Cherry

To gain insights into the genetic mechanisms regulating blooming in flowering cherry, we focused on genes categorized in four functional categories (Figure 2, Supplementary Table S3): 1) transcription factor genes; 2) phytohormone-related genes; 3) transporter and aquaporin genes; and 4) cell wall-related genes.

#### 3.4.1 Transcription factor genes

Genes encoding three types of transcription factors that trigger blooming were predominant during the flowering period. The *MYB* transcription factor genes were overrepresented from 4 to 3 WBF, while the ethylene-responsive transcription factor (*ERF*) genes and *NAC* transcription factor genes were expressed at 4 and 3 WBF, respectively.

#### 3.4.2 Phytohormone-related genes

Genes involved in biosynthesis and signal transduction pathways of gibberellin, ethylene, and cytokinin were enriched at 3–5 WBF, 3–4 WBF, and 3 WBF, respectively. Auxin-related genes were, on the other hand, expressed at 1 WBF and at the flowering days.

#### 3.4.3 Transporter and aquaporin genes

Genes encoding sugar and inorganic transporters and aquaporins that affect the turgor and osmotic pressure of cells and vacuoles were constitutively expressed from 5 WBF to the day for flowering. Among these, genes encoding 10 types of transporters (ABC transporter G family members, bidirectional sugar transporters, cationic amino acid transporters, lysine histidine transporters, organic cation/carnitine transporters, phosphate transporters, potassium transporters, UDP-galactose transporters, vacuolar amino acid transporters, and zinc transporters) were overrepresented at 3 WBF. Aquaporin genes were overrepresented at 1–2 WAF.

#### 3.4.4 Cell wall-related genes

Cell wall organization followed by cell expansions actuate petal movement, leading to flower opening. Genes encoding polysaccharide biosynthesis enzymes including UDP-glycosyltransferases and cellulose synthases were expressed from 5 WBF to the day for flowering. Genes encoding fasciclin-like arabinogalactan proteins were mainly expressed at the initial stage of the flowering period, while those encoding arabinogalactan peptides were expressed at the flowering days. Pectinesterase genes were overrepresented at the flowering days.

## 4 Discussion

Time-course RNA-seq analysis revealed the dynamics of the transcriptome of floral buds and flowers of the flowering cherry cultivar ‘Somei-Yoshino’. Subsequent WGCNA of the RNA-seq data indicated that 1,541 genes belonging to seven modules were involved in blooming at stages from 4–5 WBF to 1–2 WAF (Figure 2). In accordance with the functional annotations of these genes, those involved in floral bud and open-flower development as well as blooming were identified in flowering cherry (Figure 3). Floral bud development potentially occurs from 4–5 WBF, with the gibberellin-mediated activation of genes controlling sugar and polysaccharide metabolism. Subsequently, at 3–4 WBF, ethylene and cytokinin likely accelerate the expression of *MYB, ERF*, and *NAC* transcription factor genes, resulting in the initiation of cell wall organization and cell division (Nakano et al., 2015). Some *MYB* and *NAC* genes are also involved in gametophyte or ovule integument development (Kunieda et al., 2008; Renak et al., 2012). According to the observations in sweet cherry (*Prunus avium*), the 3–4 WBF timepoint corresponds to the formation of tetrads or microspores after meiosis in the pollen (Fadon et al., 2019). At this stage, amino acid metabolism is also upregulated. Metabolome analysis of the flower buds of Japanese apricot (*Prunus mume*) revealed significant changes in the abundance of several amino acids before the bud break stage (Zhuang et al., 2015). Amino acids play important roles in bud break. At 1–2 WBF, the reproduction process, together with metabolism, is likely activated. Auxin-related genes were also activated at this stage in this current study. On the flowering day, we speculate that the auxin signal alters pectinesterase gene expression to initiate cell wall remodeling, leading to rapid petal enlargement and movement, which is the most dynamic movement in flowering plants (Habu and Tao, 2014; Nakano et al., 2015). The high-level expression of aquaporin genes, which encode water channel proteins localized to the plasma membrane or tonoplast membranes, on the day of flowering might facilitate the water flux into the petals to allow smooth petal movement (Azad et al., 2013). The relationship between auxin signaling and pectinesterase gene expression has been demonstrated in the ripening process of strawberry fruit (Castillejo et al., 2004). Similar hormonal regulations might be conserved in the Rosaceae species. The role of the auxin signal in the dynamic movement of petals during flower opening needs further investigation. After blooming, genes involved in the stimulus response and cell-to-cell communication are expressed the flower, especially in the pistil and ovary, which might reflect the pollination and fertilization process, even though ‘Somei-Yoshino’ is self-incompatible. Similar transcriptome profiles were observed in other *Prunus* species (Habu and Tao, 2014; Gómez et al., 2019; Iqbal et al., 2021).

**Figure 3.**
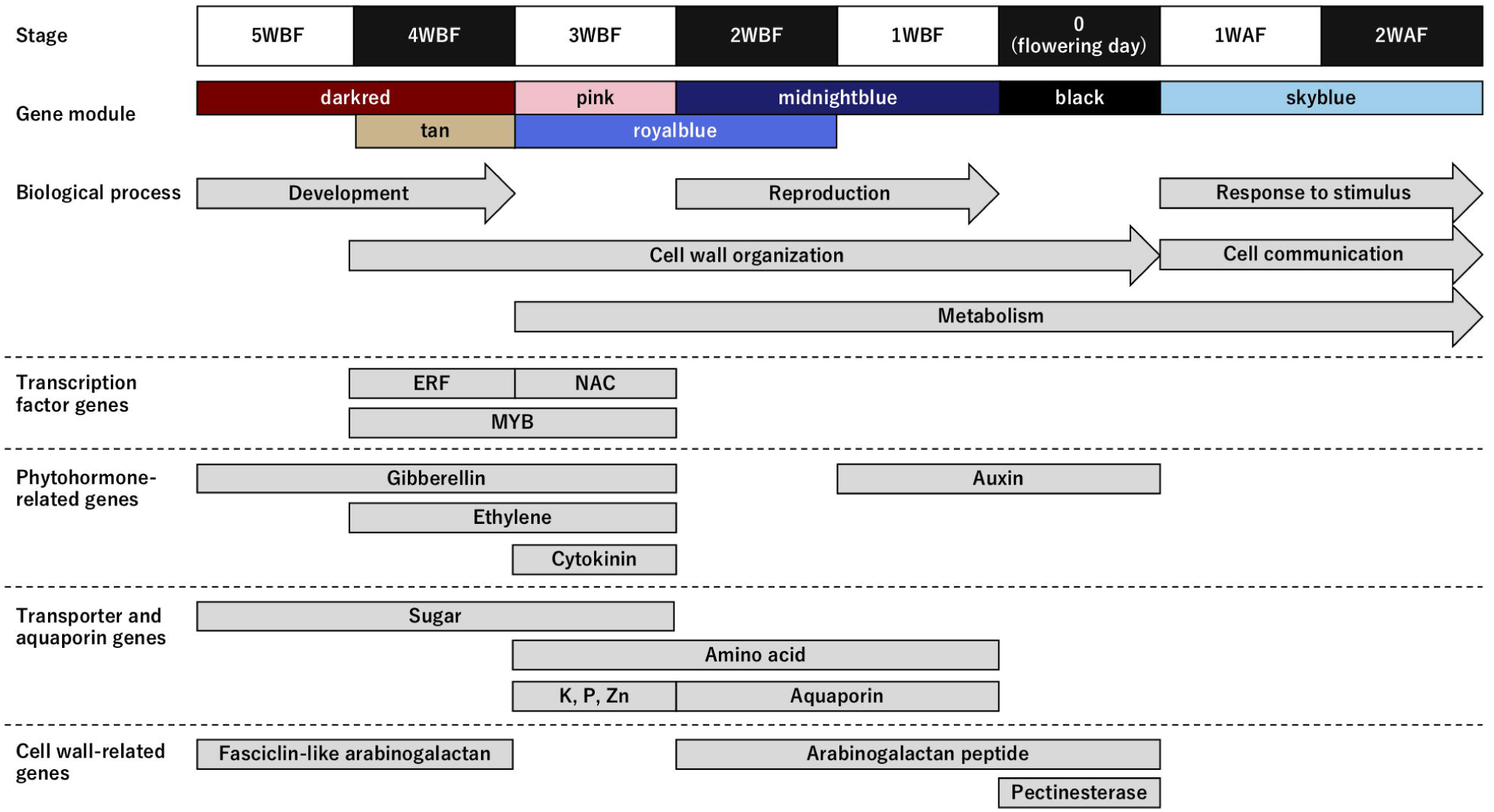
Genes enriched in the biological process (BP) category and involved in the flower opening mechanism in ‘Somei-Yoshino’. Upper panels show the time frame (from 5 weeks before flowering [WBF] to 2 weeks after flowering [WAF]) and the corresponding gene modules. Arrows indicate the Gene Ontology (GO) terms enriched in the BP category (see Table 1). Boxes indicate the properties of the genes highly expressed in the seven characterized modules (see Supplementary Table S3).

Because the flowering time of ‘Somei-Yoshino’ is important for the tourism industry in Japan, methods for predicting the flowering date of this cultivar have been developed in accordance with the cumulative temperature before flower opening (Aono and Murakami, 2017). Gene expression analysis could upgrade the current forecasting model, for which genes expressed on specific days before flowering could be employed as diagnostic markers. However, practical RNA quantification methods, such as real-time quantitative PCR and the next-generation sequencing (NGS)-based RNA-seq, are time-consuming and labor-intensive. Since diagnostic markers can be detected easily within a short time, easy-to-use target-RNA qualifying methods are required to predict the flowering time in ‘Somei-Yoshino’, based on gene expression data.

Comparative time-course transcriptome analysis would enable the selection of high-confidence diagnostic markers for various purposes. For example, time-course transcriptome analysis can be applied to floral buds of vegetable and fruit crops and cut flowers. Moreover, this comparative analysis is not only limited to the selection of markers for forecasting flowering time but is also applicable to the prediction of disease, fertilization, and harvest time in the field. Indeed, transcriptome profiling has been employed to monitor nutritional responses and adaptation in rice (Takehisa and Sato, 2021).

Overall, this study provides insights into the genetic mechanisms controlling petal movement and blooming in cherry. Further studies are required to connect the genetic insights with the physiological mechanisms (van Doorn and Van Meeteren, 2003). Once the connection or relationship is validated, the transcriptome-based prediction would serve as a powerful tool for monitoring the plant phenotype under controlled cultivation conditions as well as in the field. For example, genome-based prediction has been used to predict progeny phenotypes in breeding programs. The transcriptome- and genome-based predictions would promise high-confidence forecasting plant traits in the near and distant future, respectively.

## 5 Conflict of Interest

The authors declare that the research was conducted in the absence of any commercial or financial relationships that could be construed as a potential conflict of interest.

## 6 Author Contributions

KS, TE, and AI conceived this study and collected the samples; KS and SI performed experiments and collected data; KS, TE, and AI analyzed and interpreted the data; KS wrote the manuscript with contributions from TE; and all authors read and approved the final manuscript.

## 7 Funding

This work was supported by the Kazusa DNA Research Institute Foundation.

## 8 Acknowledgments

We thank Y. Kishida, C. Minami, S. Sasamoto, H. Tsuruoka, and A. Watanabe (Kazusa DNA Research Institute) for providing technical assistance.

## Notes

### Competing Interest Statement

The authors have declared no competing interest.

