## Supplementary Information for "Transcriptome Dynamics of Floral Organs Approaching Blooming in the Flowering Cherry (*Cerasus* × *yedoensis*) Cultivar ‘Somei-Yoshino’"

**Supplementary Table S1** Sampling dates of 'Somei-Yoshino' floral buds and flowers.

**Supplementary Table S2** Gene modules overrepresented at different flowering stages.

**Supplementary Table S3** Genes and annotations overrepresented in the seven modules.

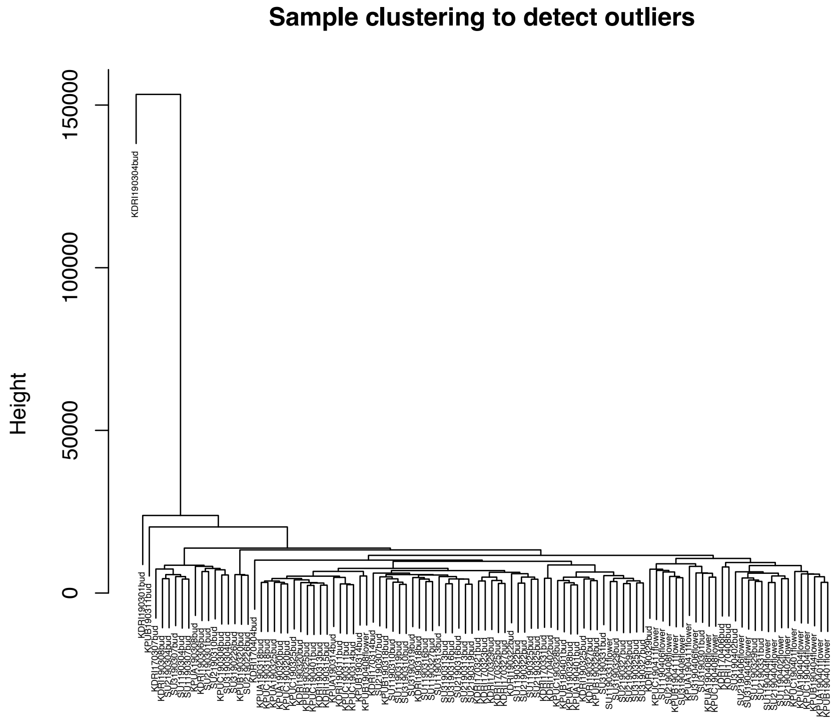


**Supplementary Figure S1** Sample clustering used to identify outliers.

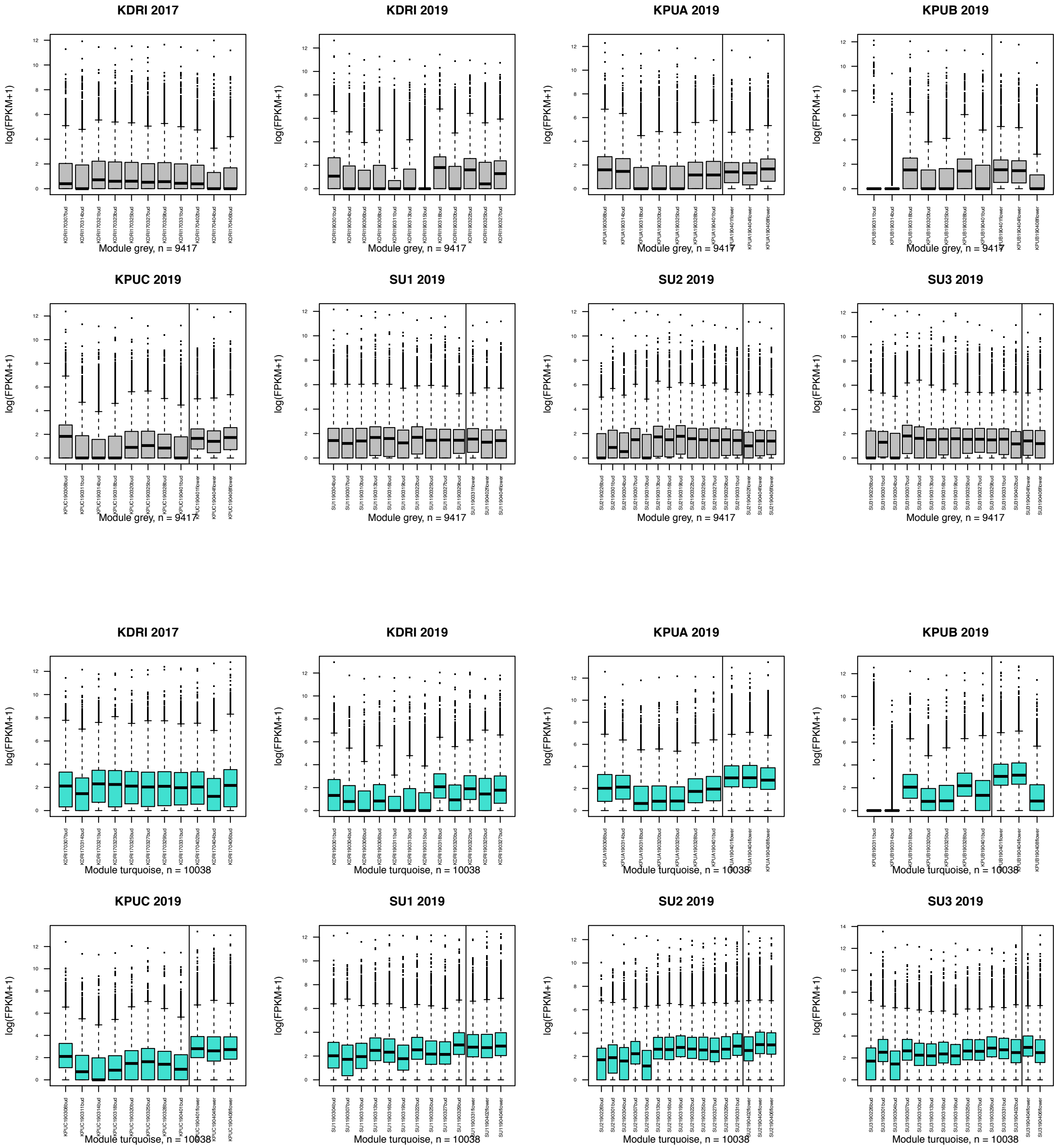


**Supplementary Figure S2** Expression patterns of gene modules, based on weighted gene correlation network analysis (WGCNA). Vertical lines indicate the flowering day.

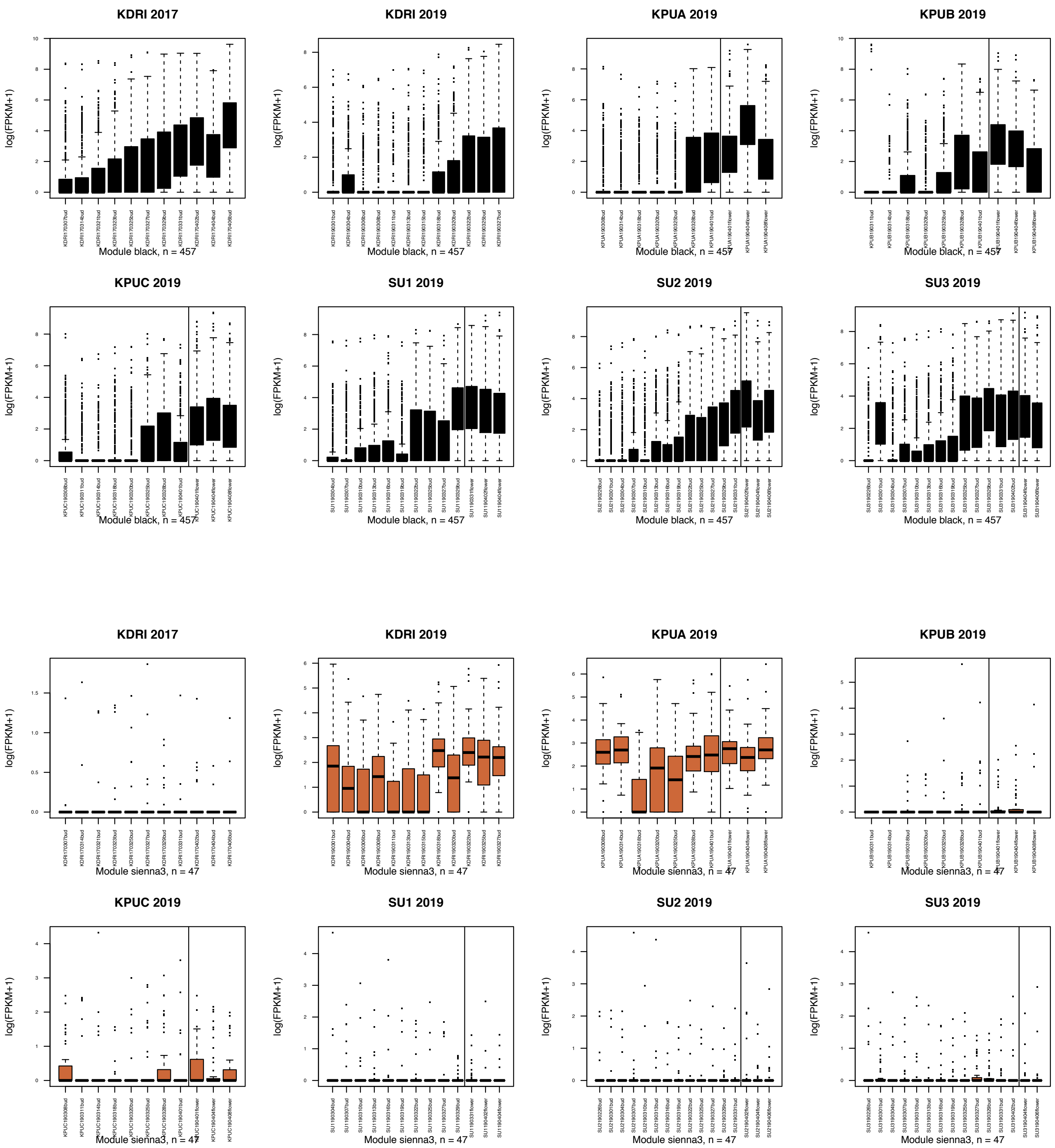


**Supplementary Figure S2** (Continued.)

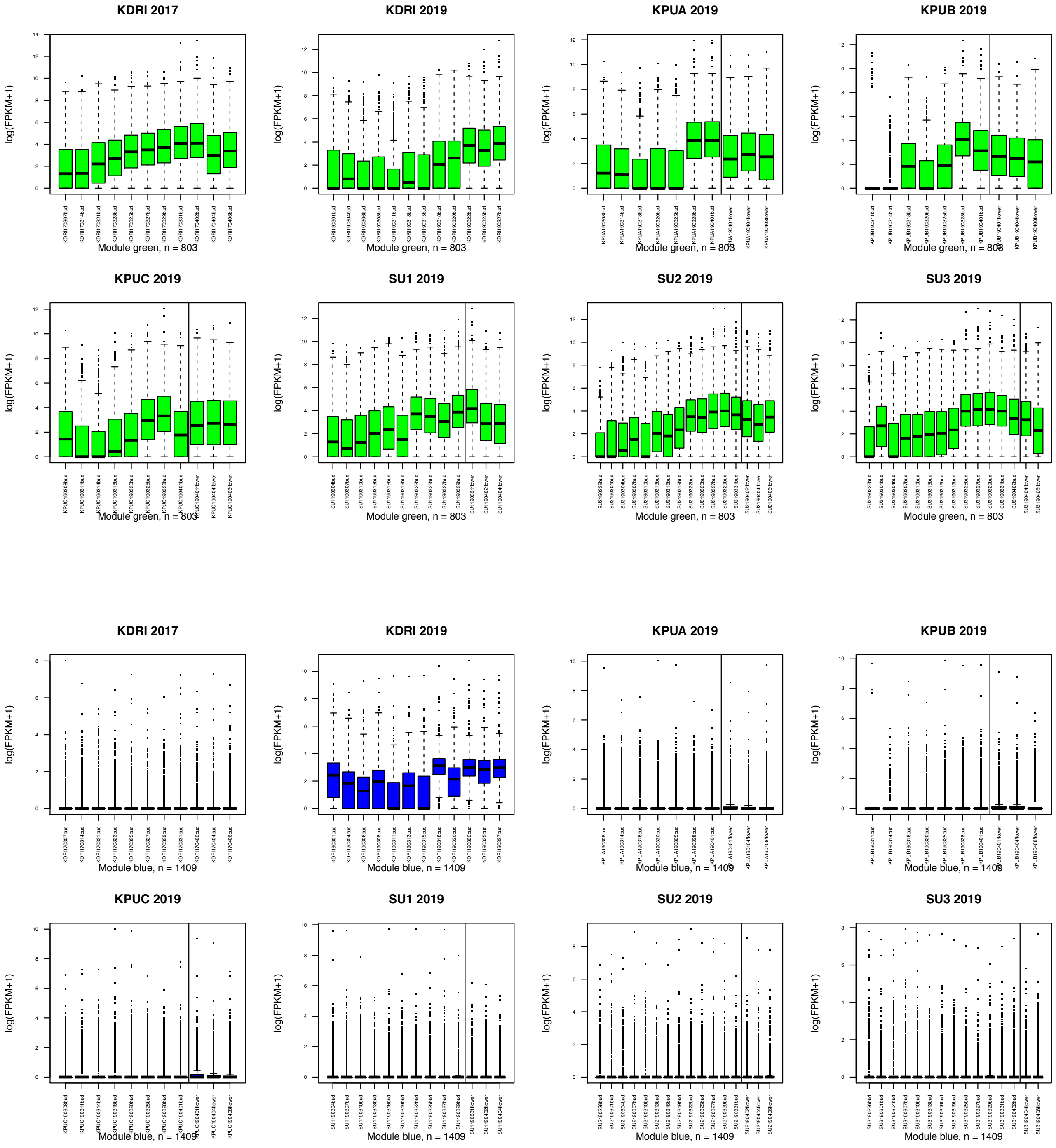


**Supplementary Figure S2** (Continued.)

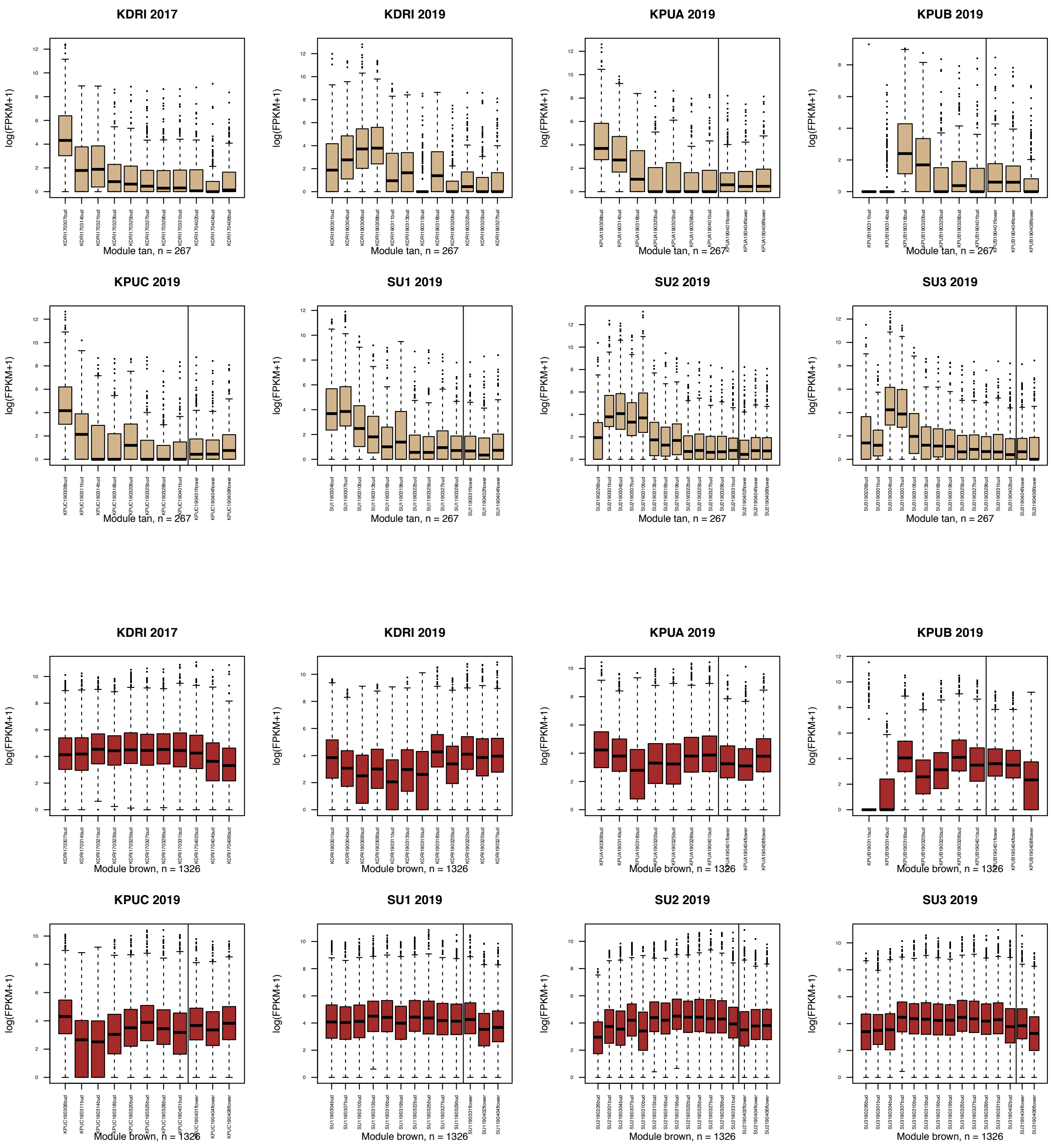


**Supplementary Figure S2** (Continued.)

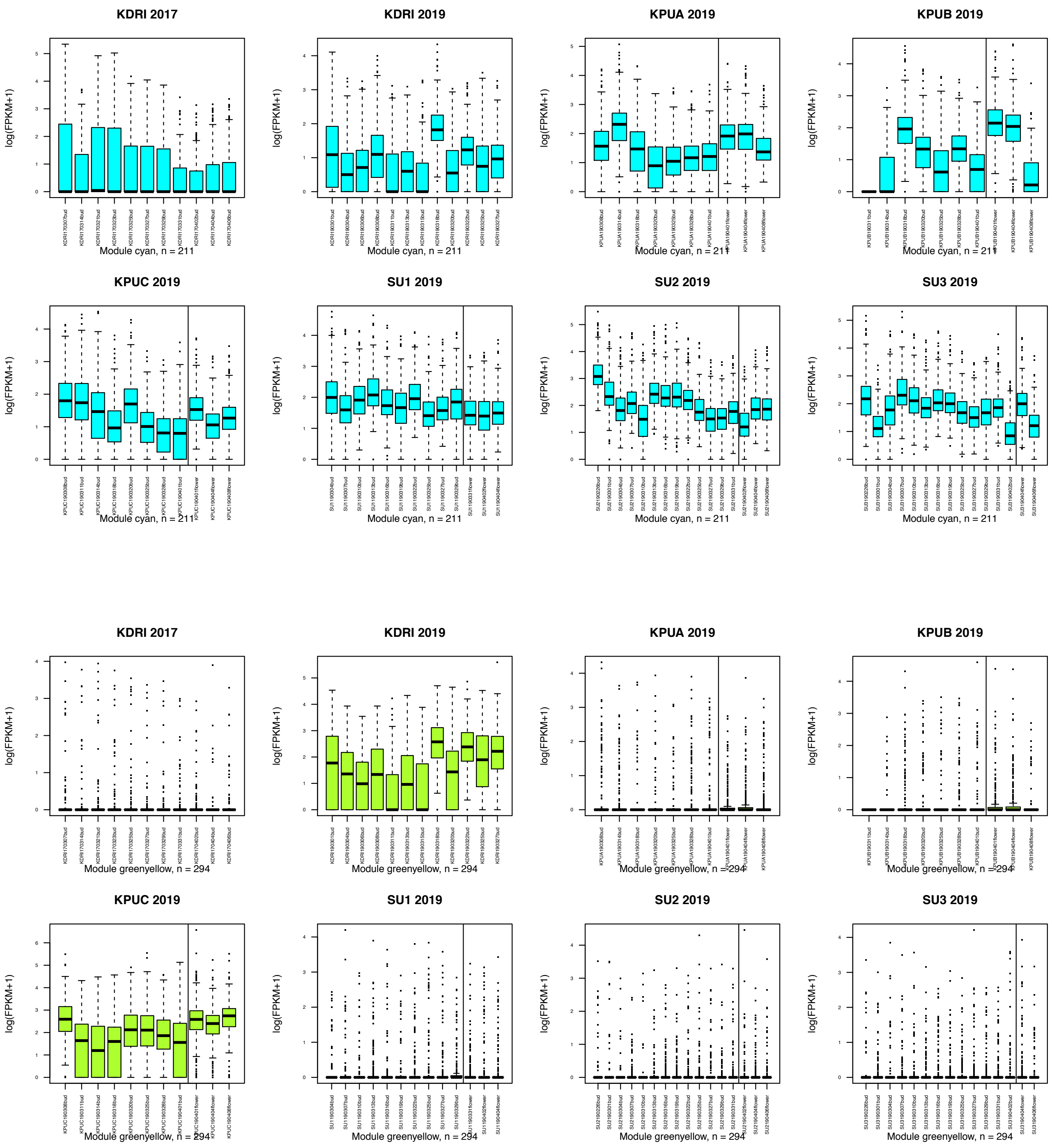


**Supplementary Figure S2** (Continued.)

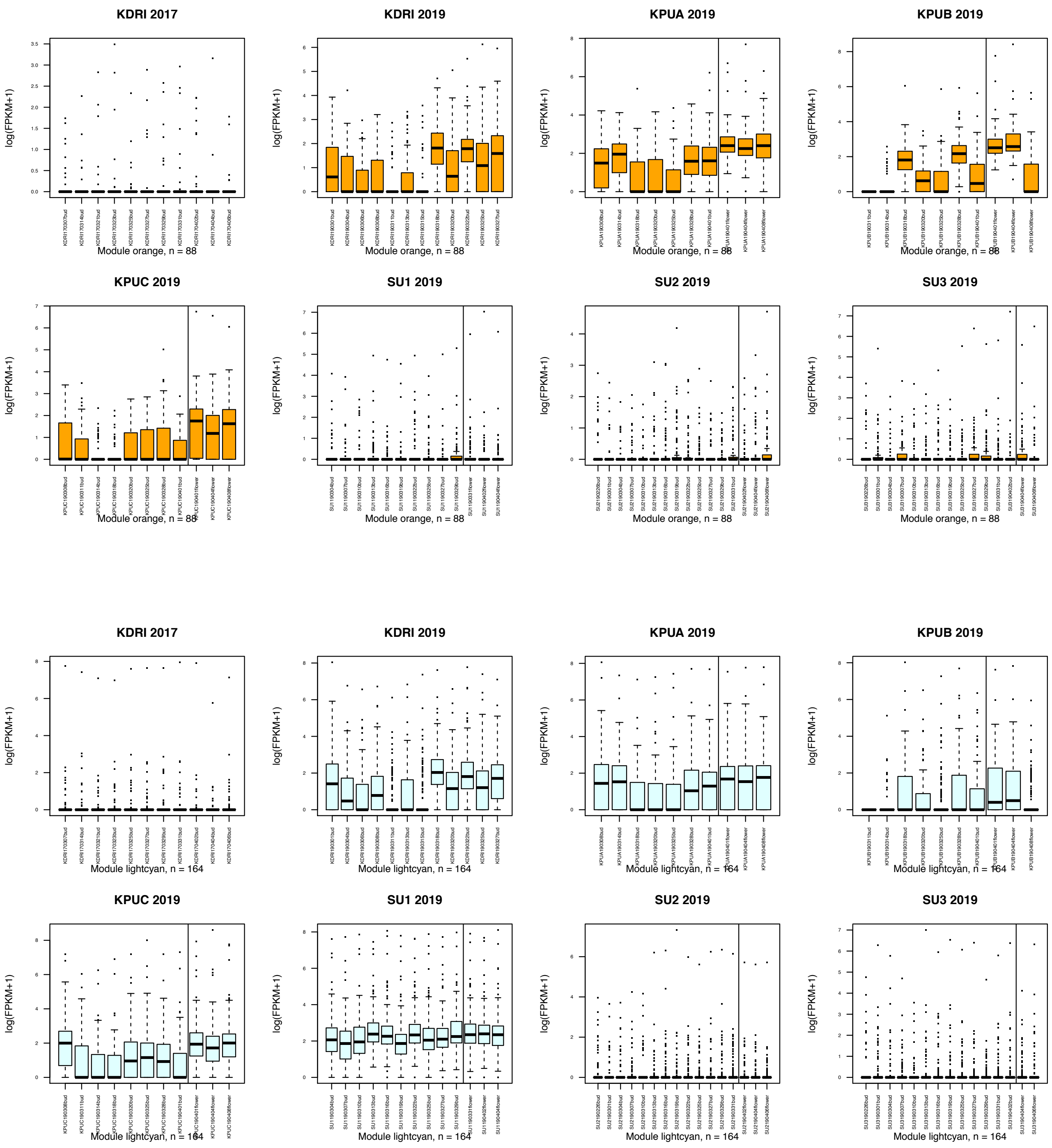


**Supplementary Figure S2** (Continued.)

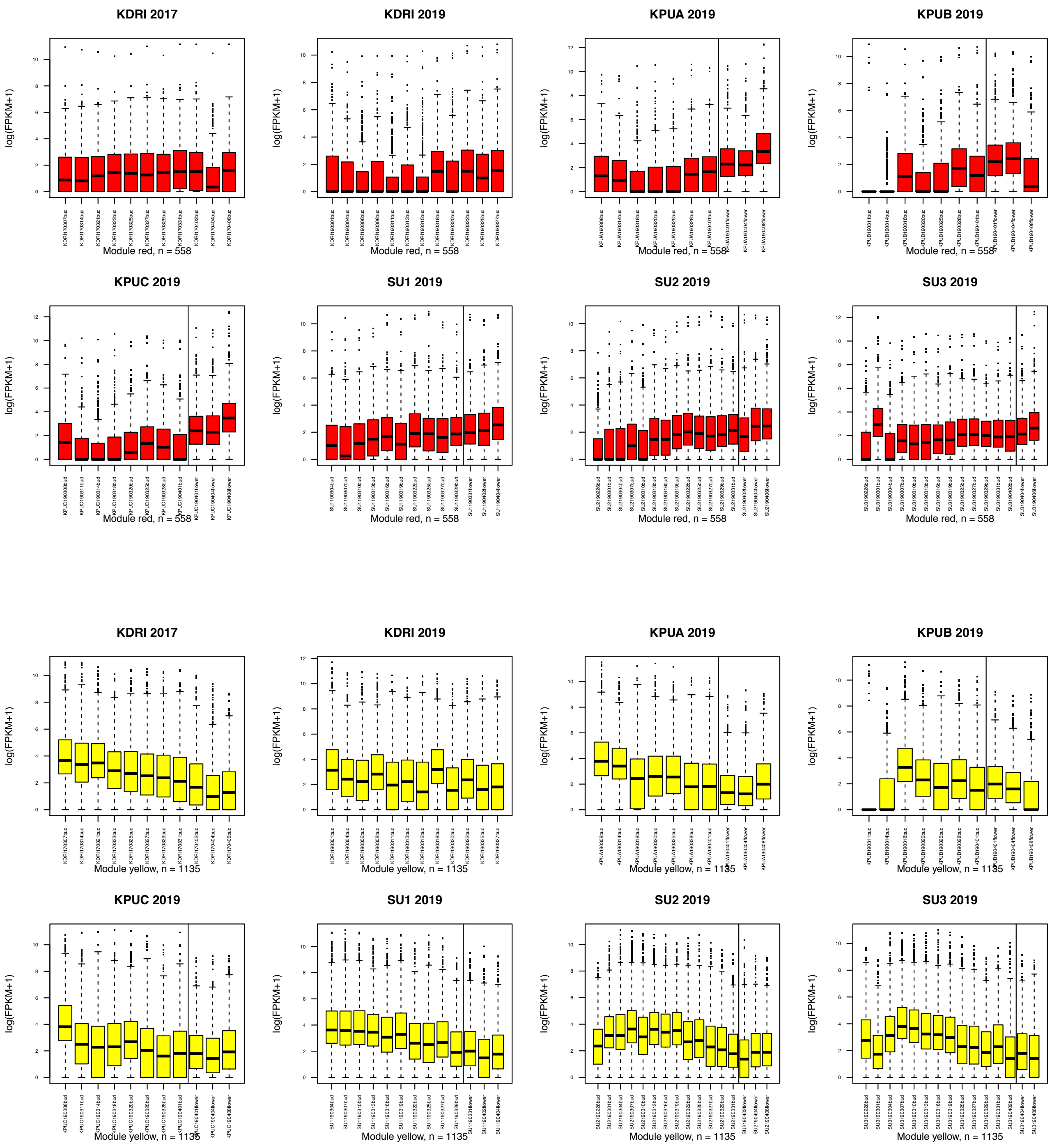


**Supplementary Figure S2** (Continued.)

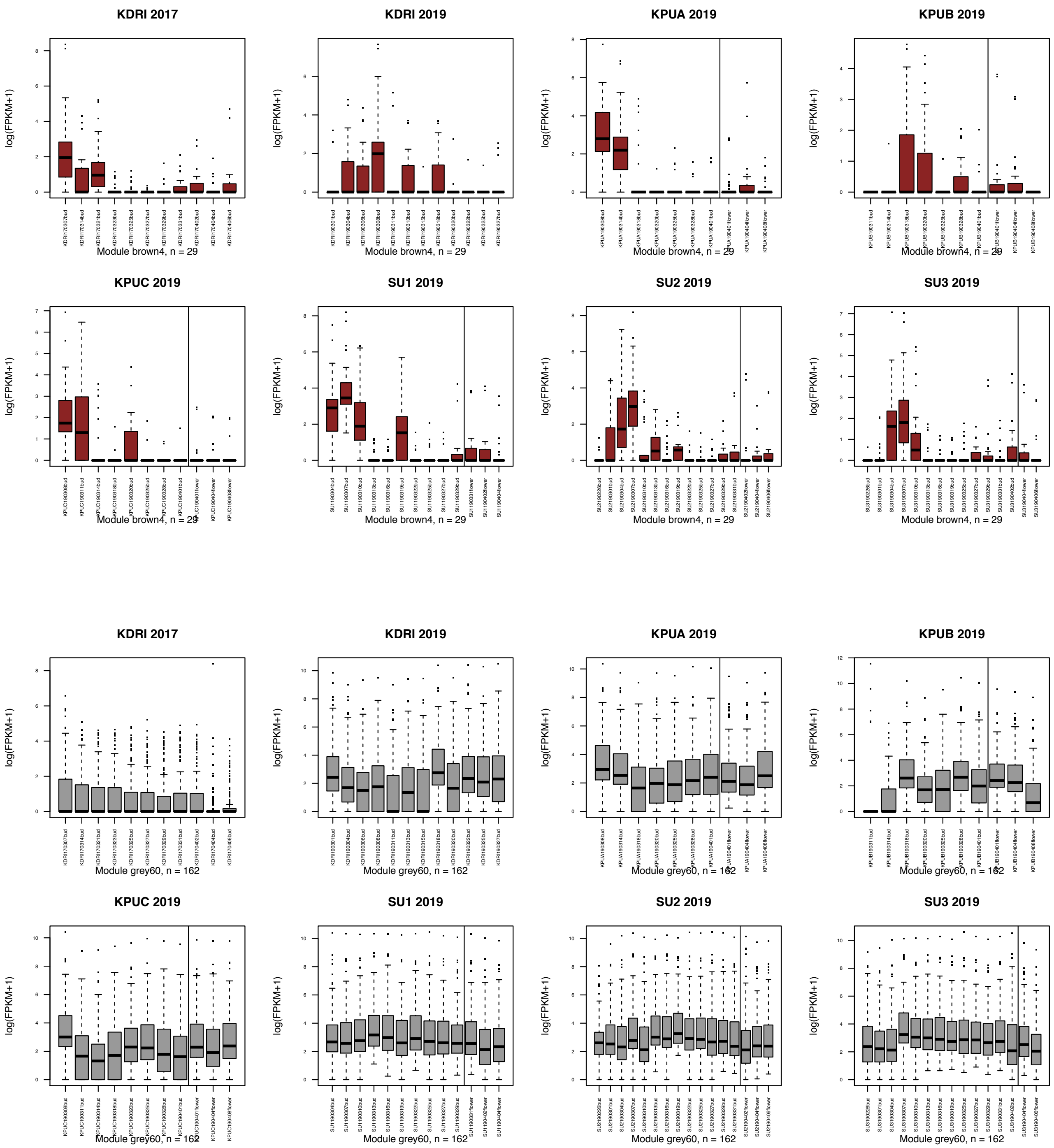


**Supplementary Figure S2** (Continued.)

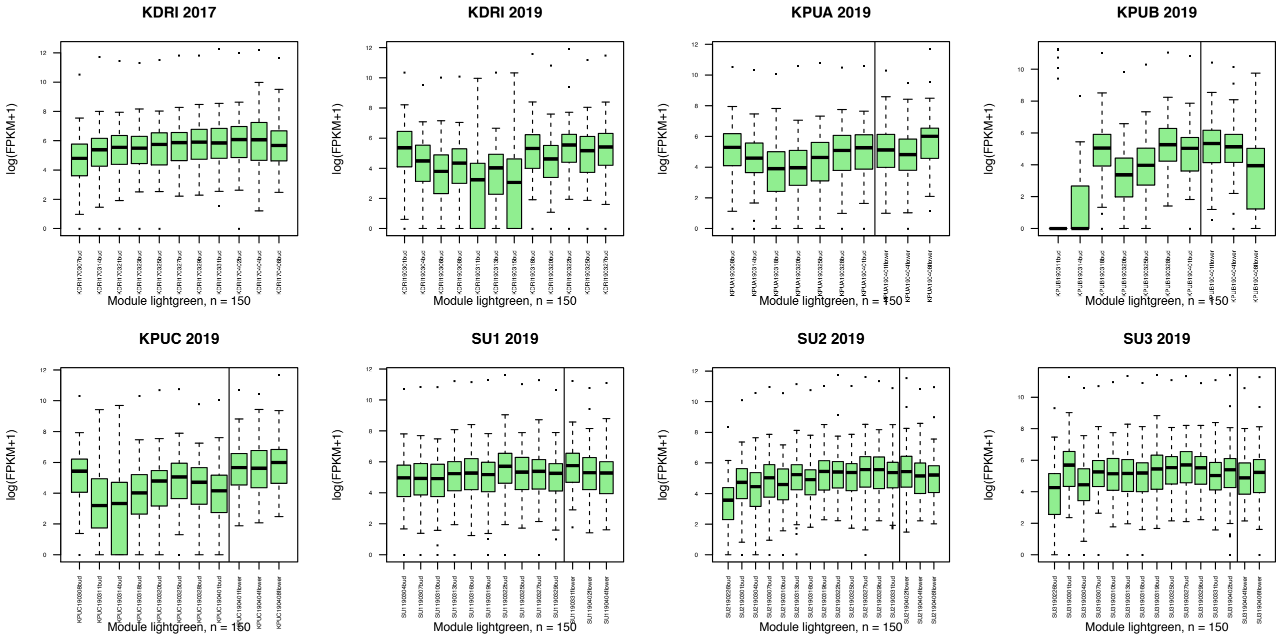


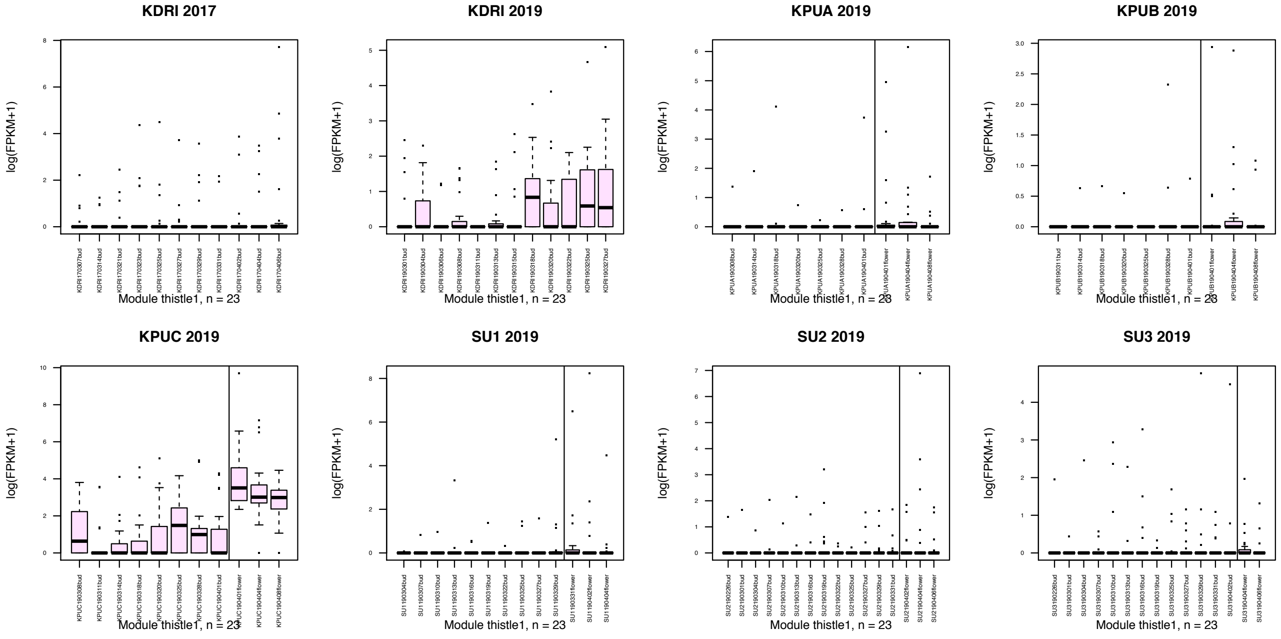


**Supplementary Figure S2** (Continued.)

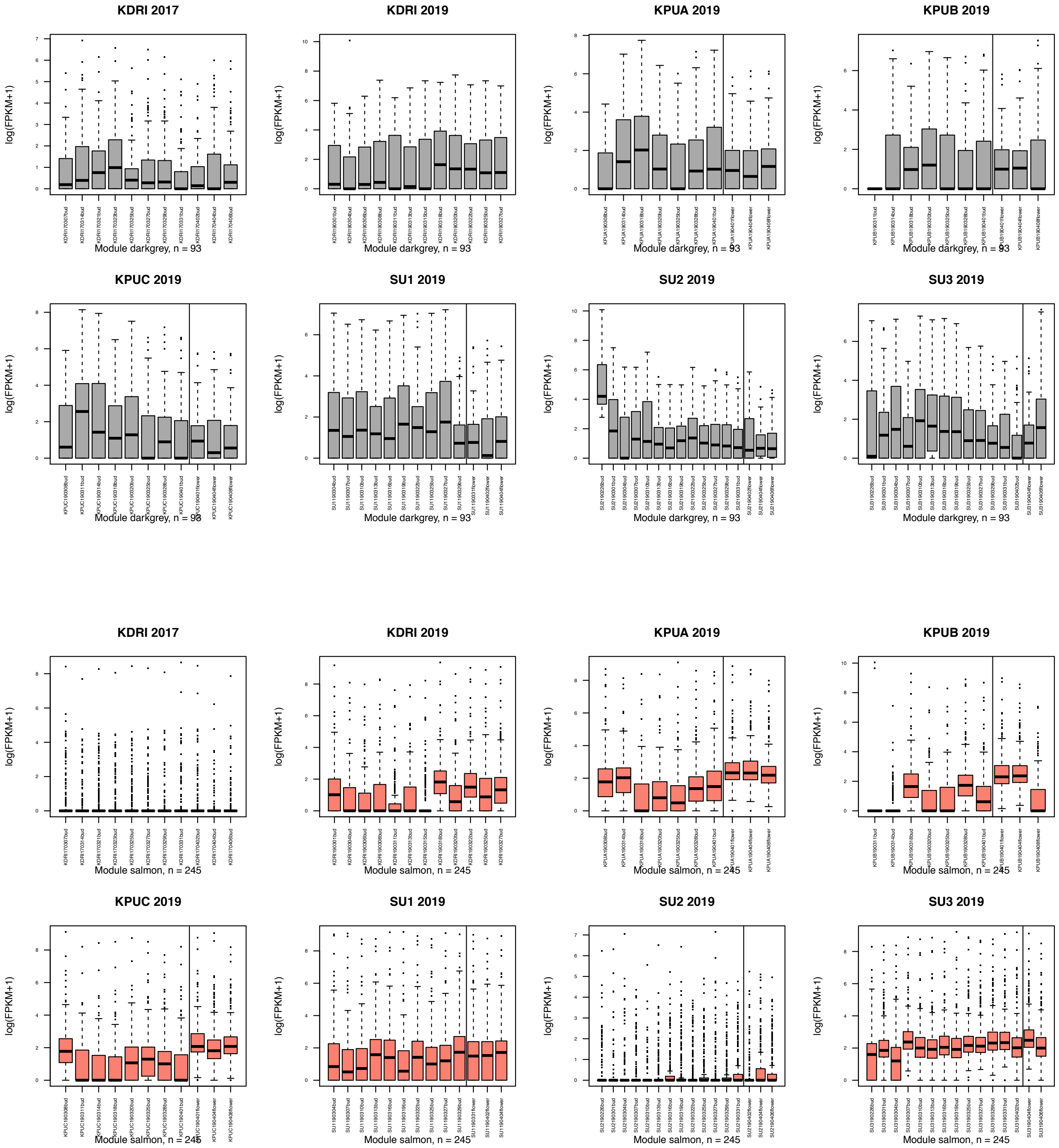


**Supplementary Figure S2** (Continued.)

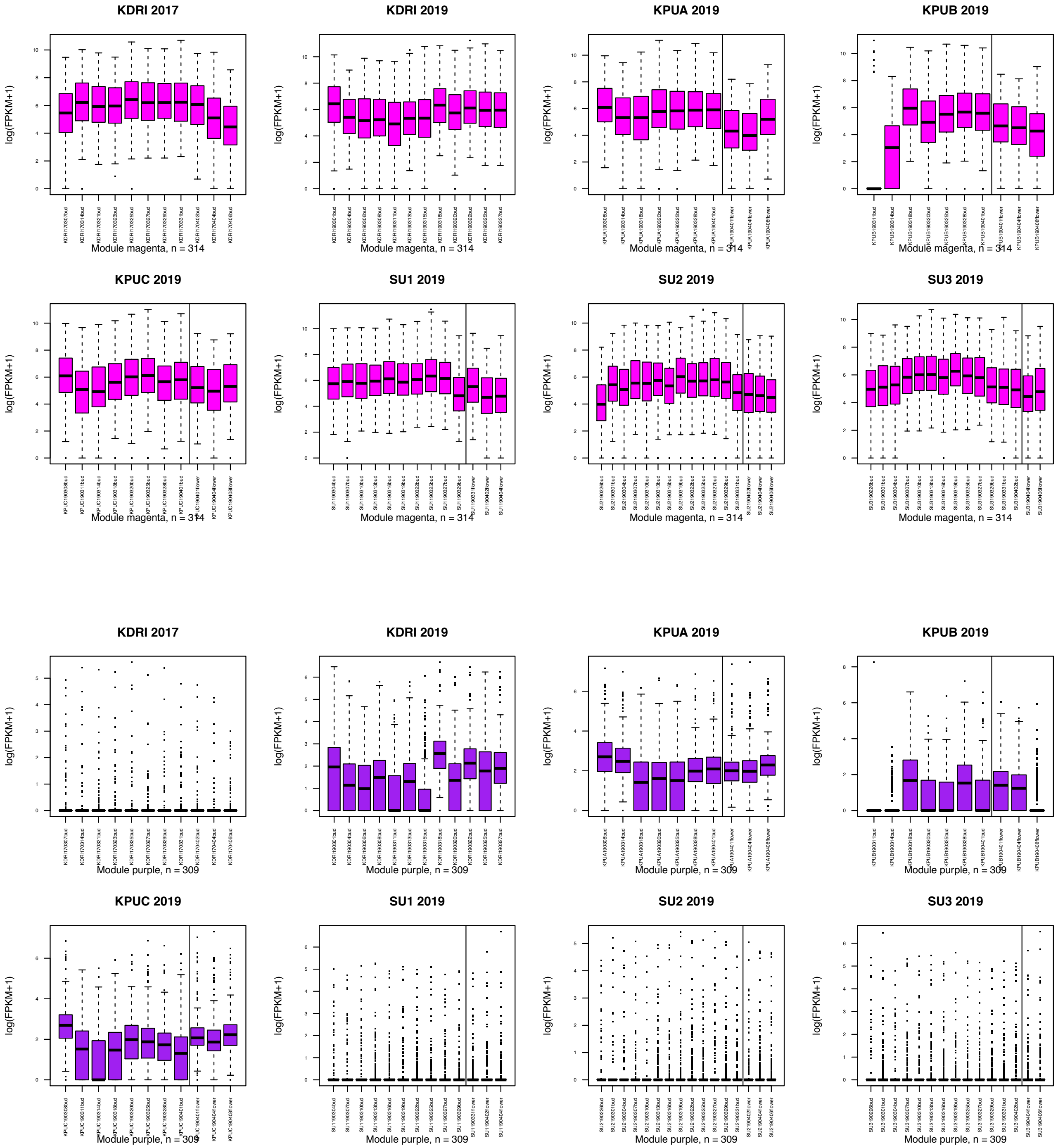


**Supplementary Figure S2** (Continued.)

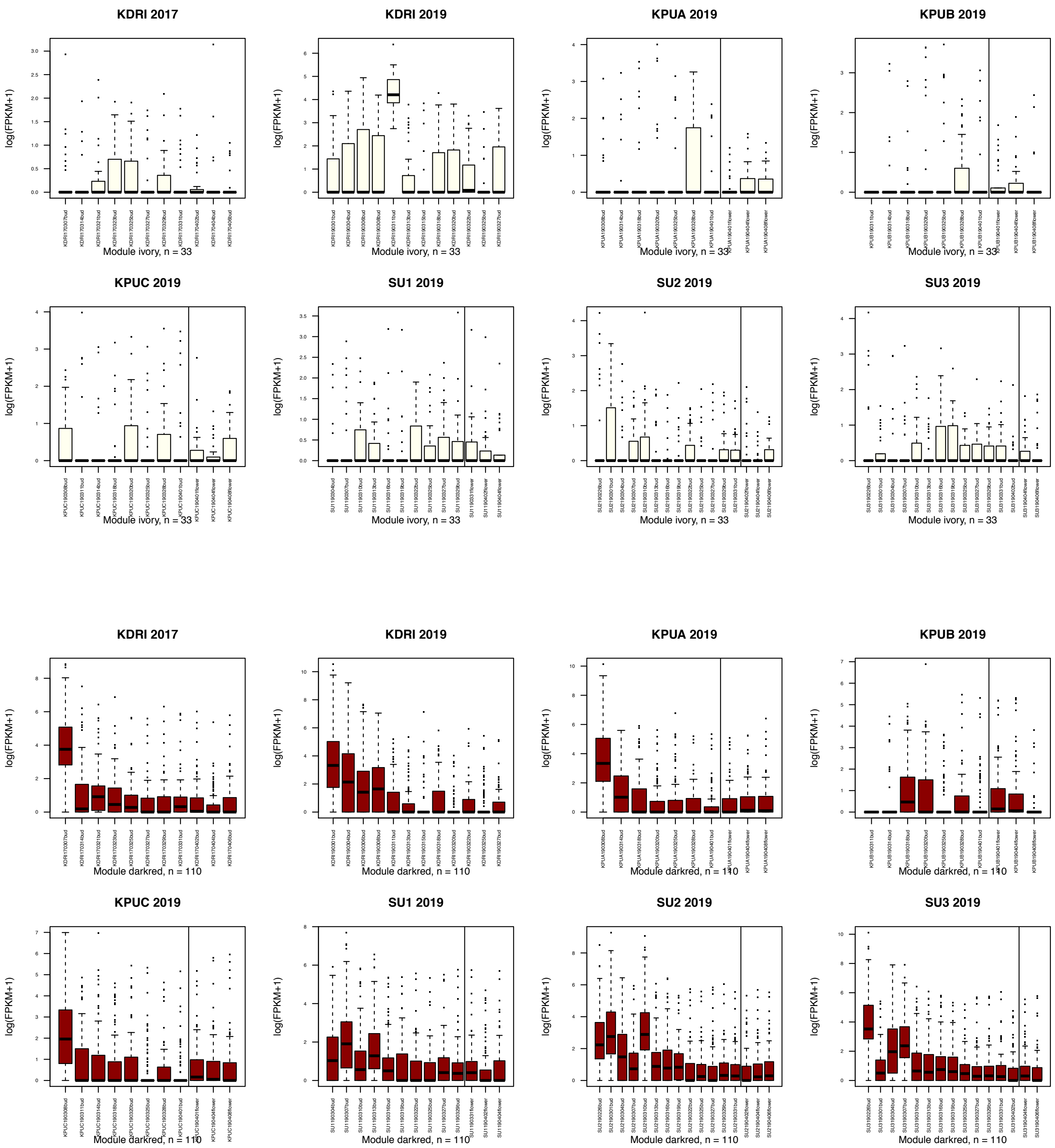


**Supplementary Figure S2** (Continued.)

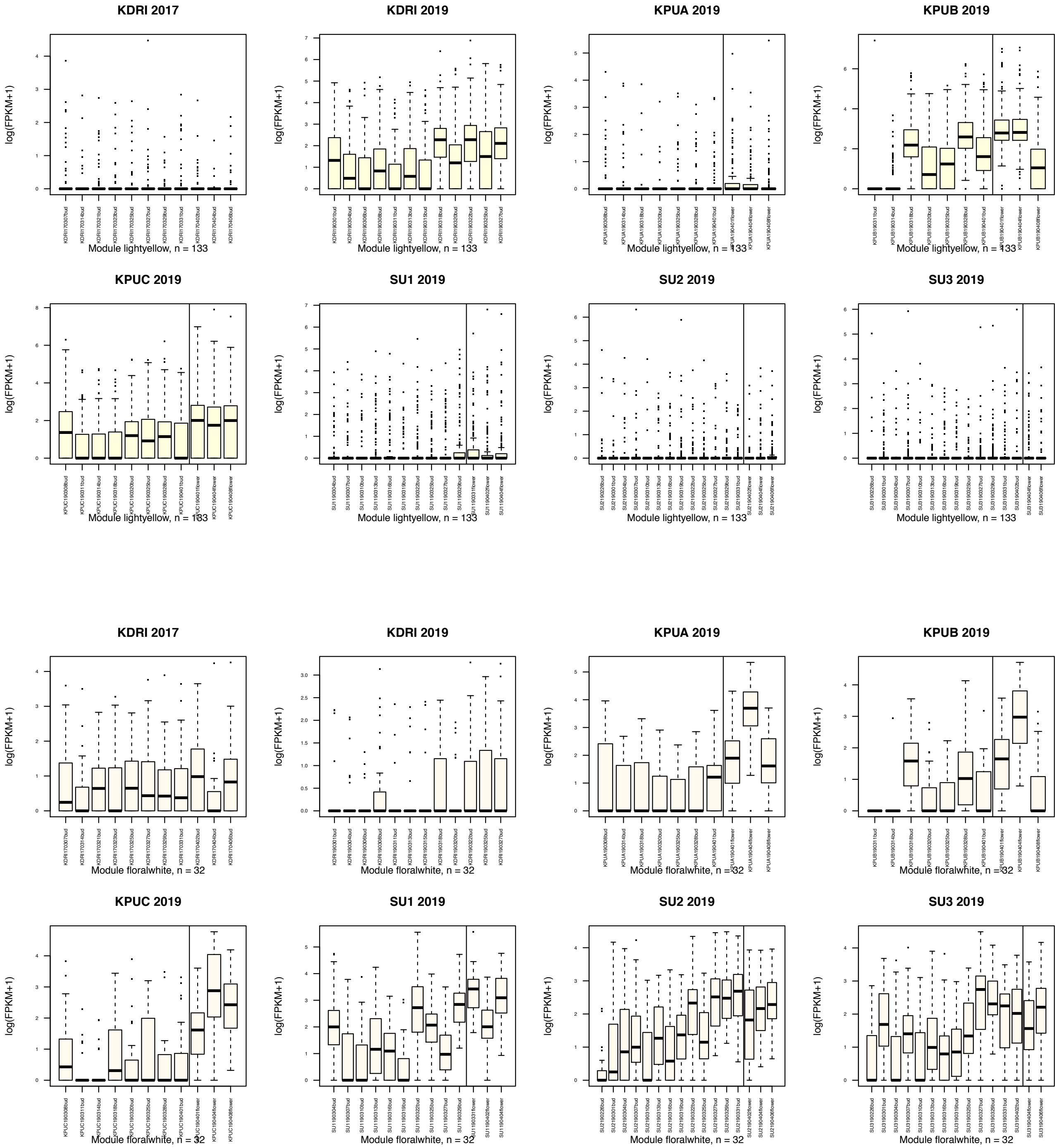


**Supplementary Figure S2** (Continued.)

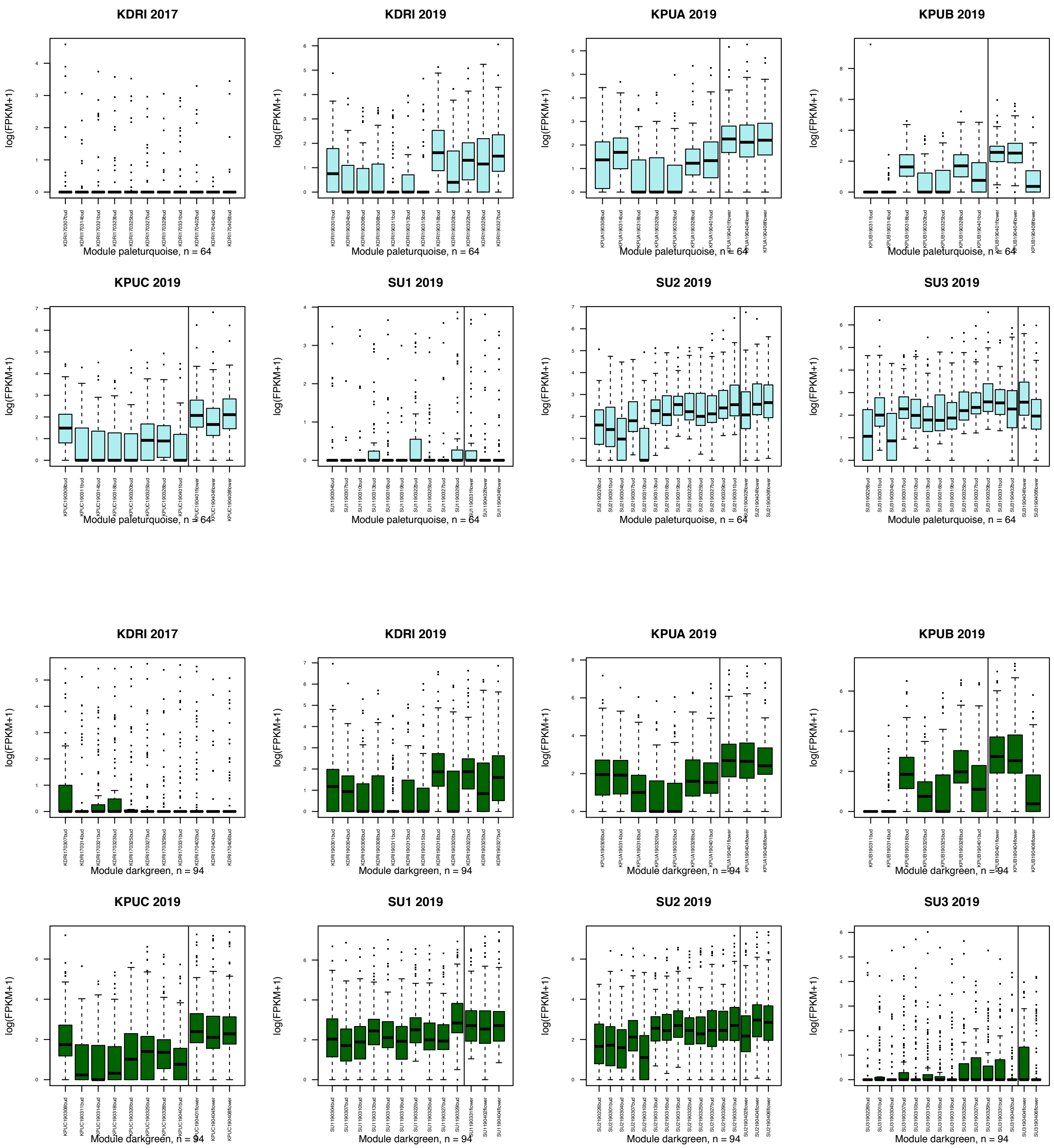


**Supplementary Figure S2** (Continued.)

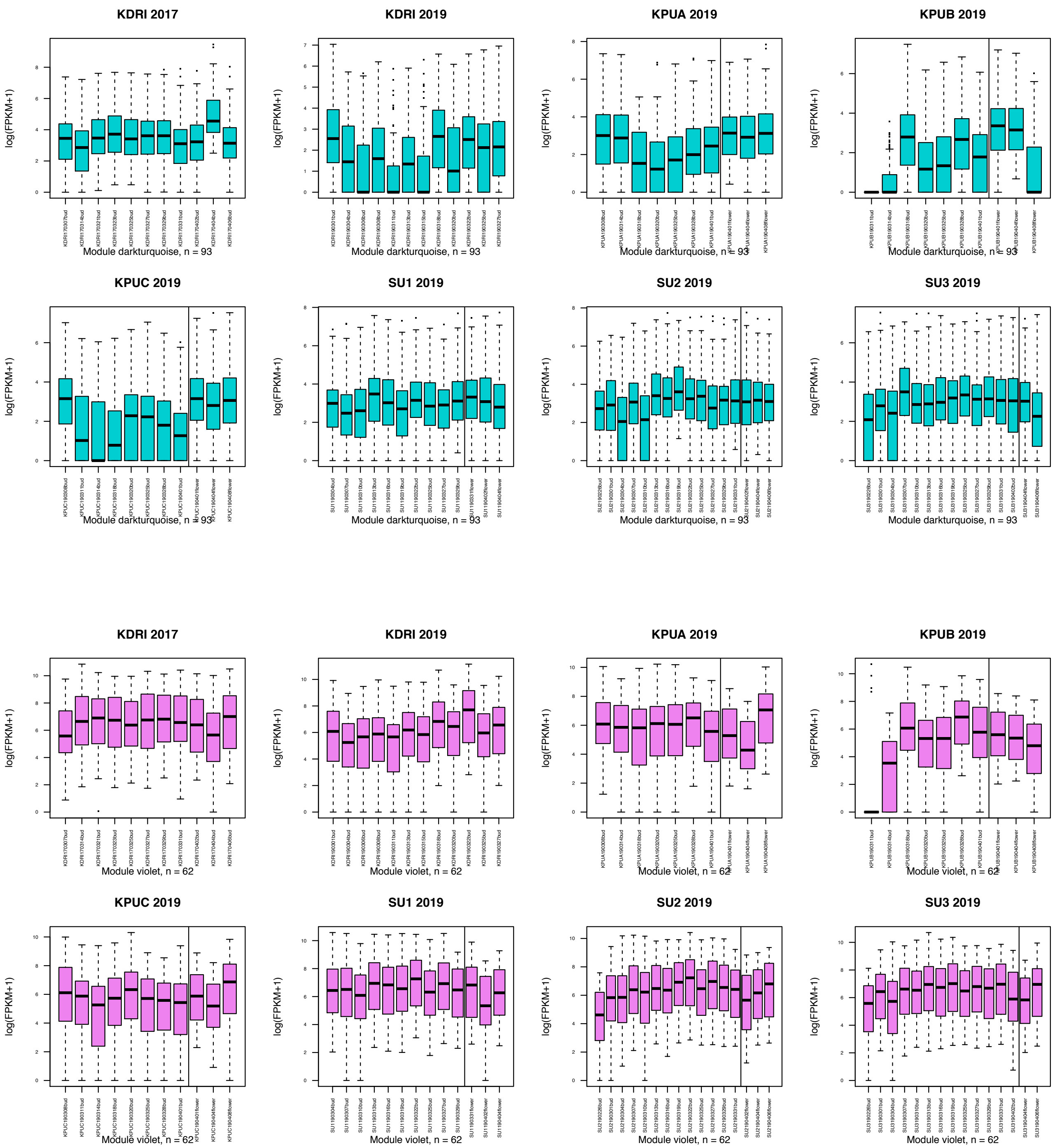


**Supplementary Figure S2** (Continued.)

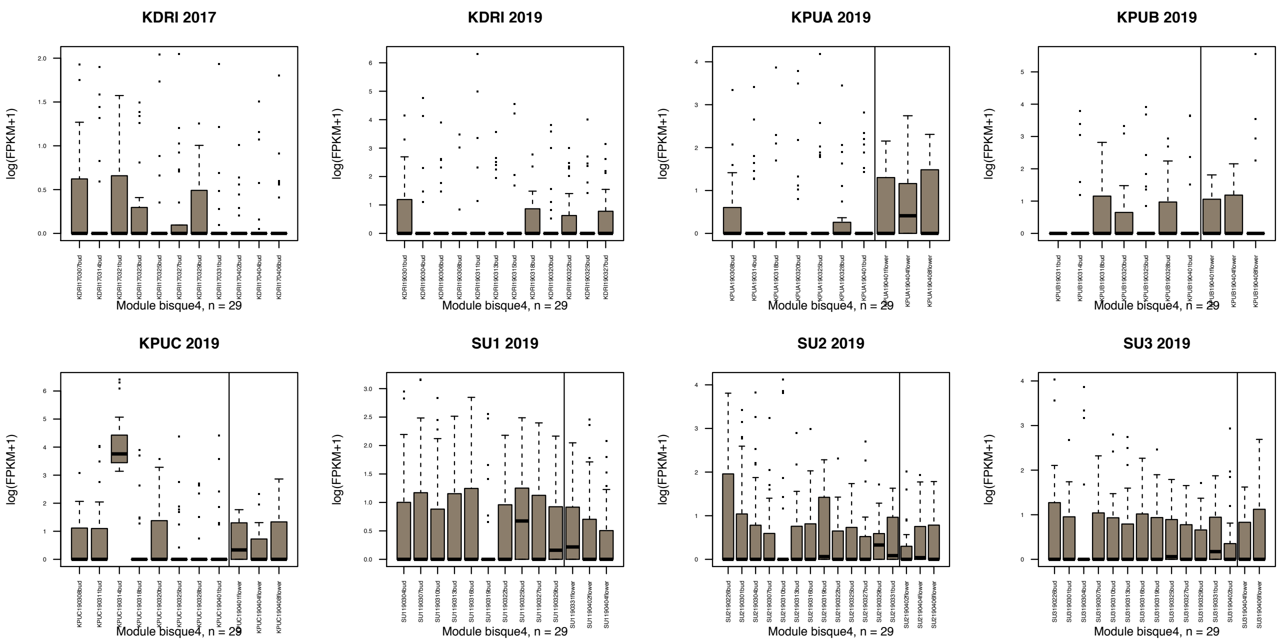


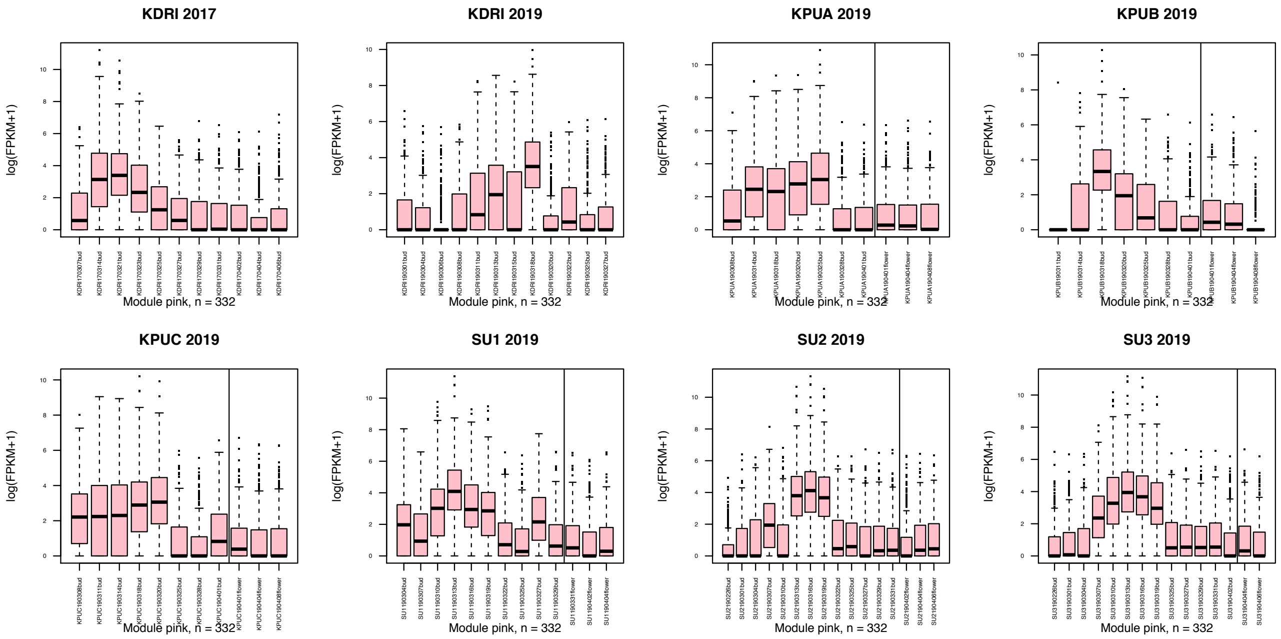


**Supplementary Figure S2** (Continued.)

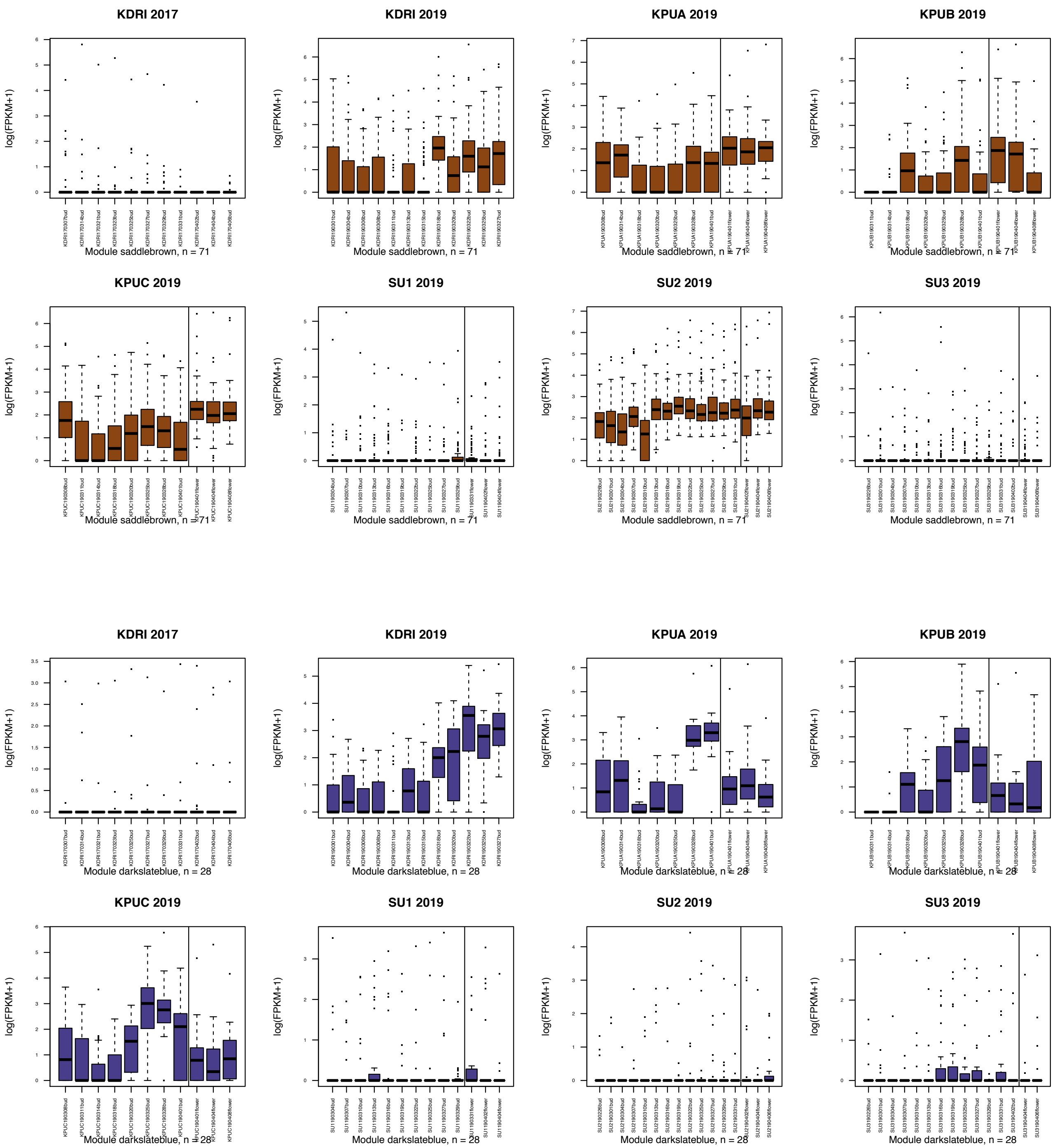


**Supplementary Figure S2** (Continued.)

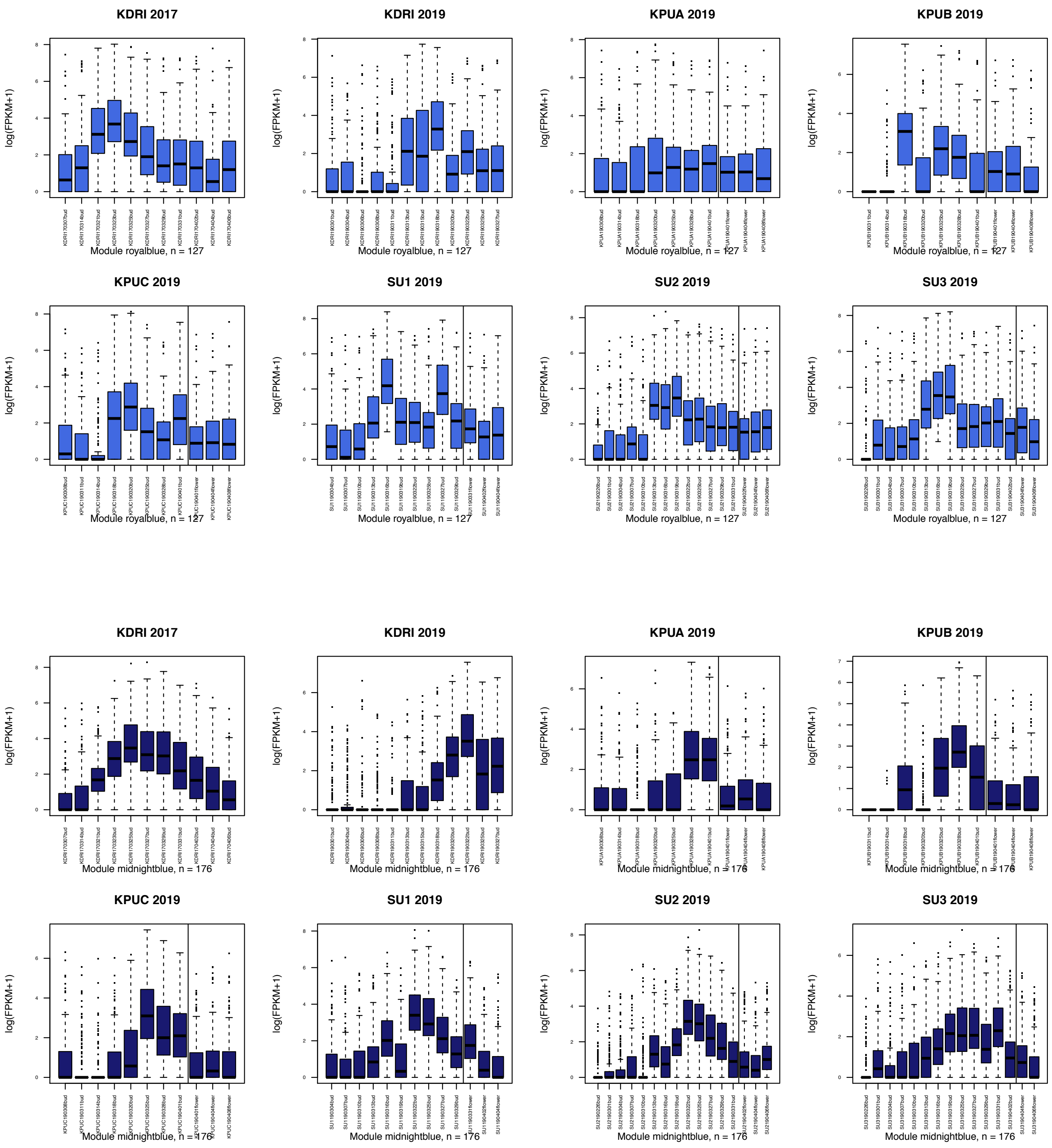


**Supplementary Figure S2** (Continued.)

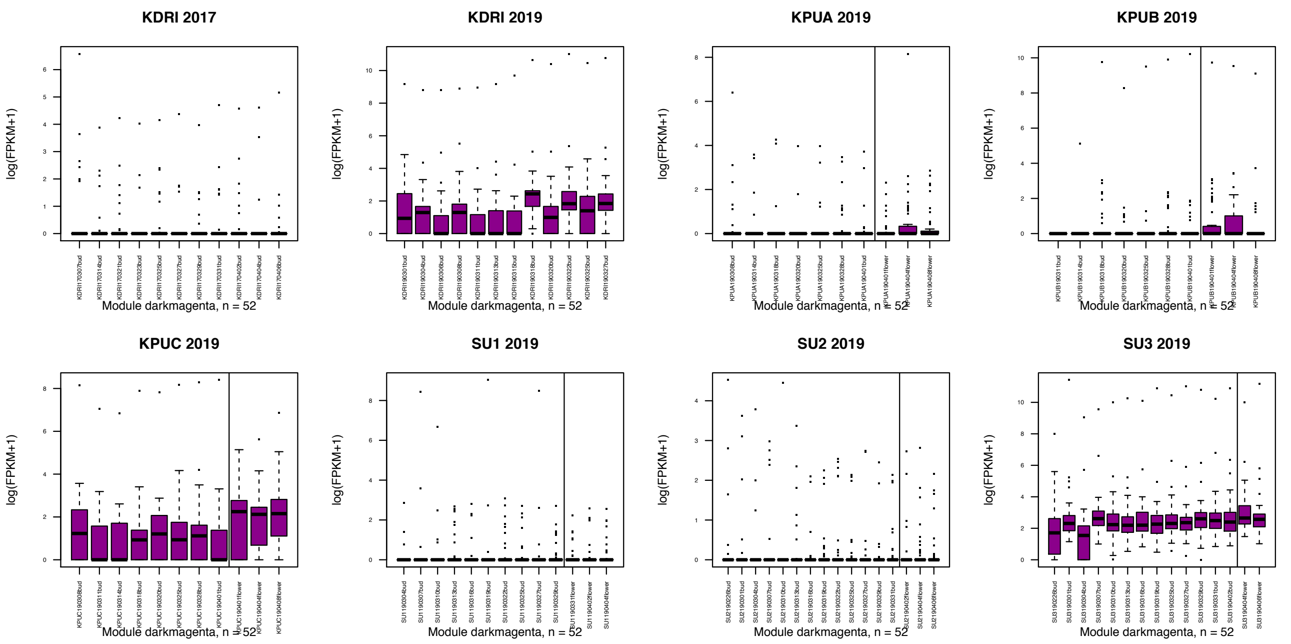


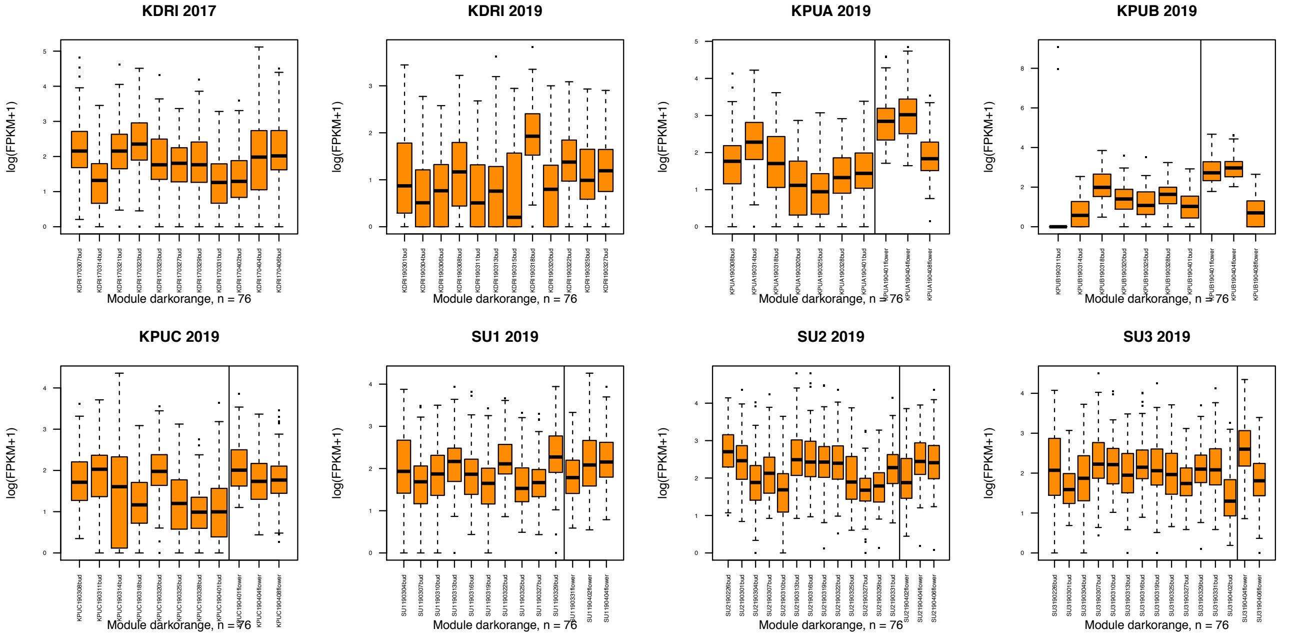


**Supplementary Figure S2** (Continued.)

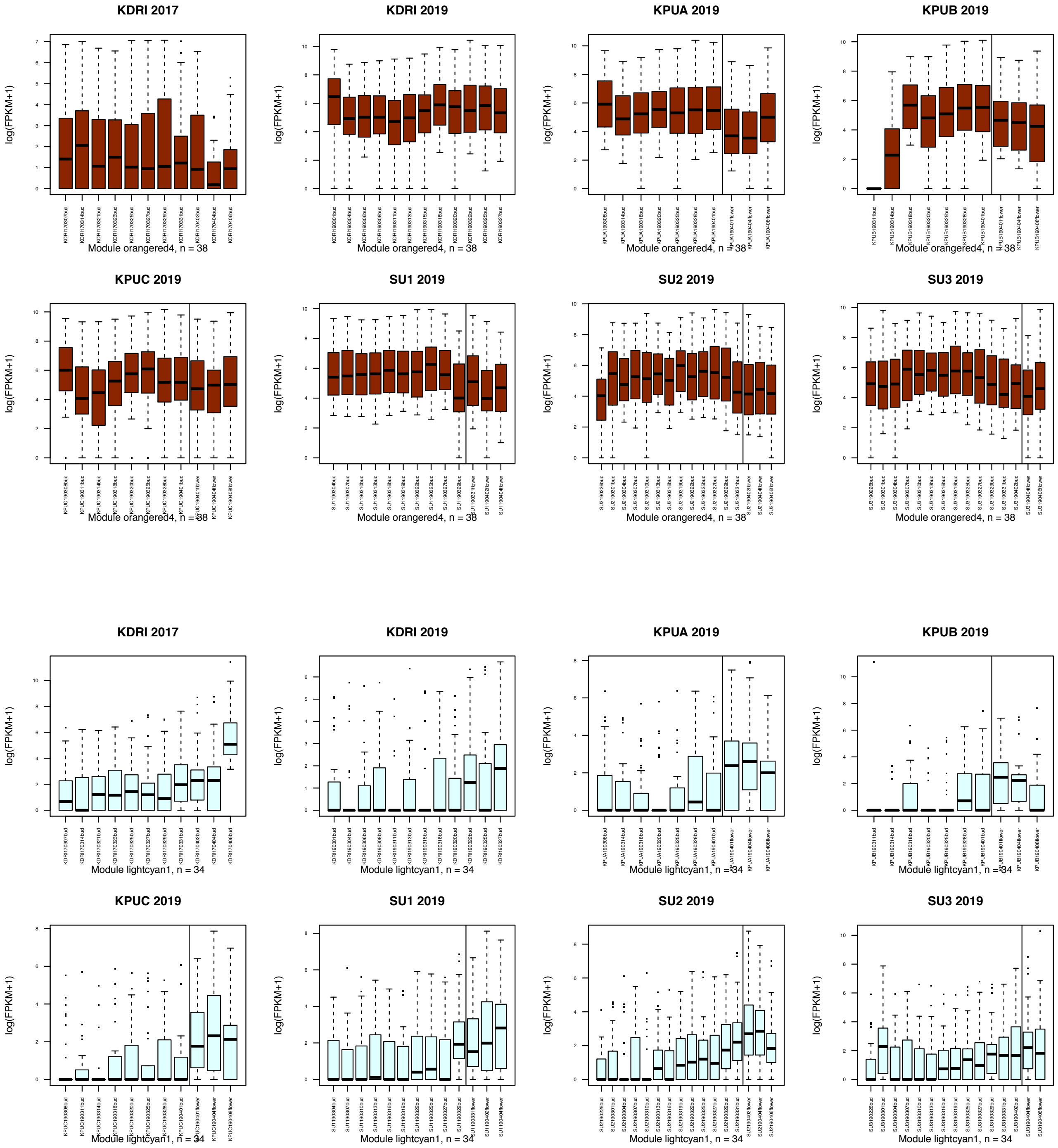


**Supplementary Figure S2** (Continued.)

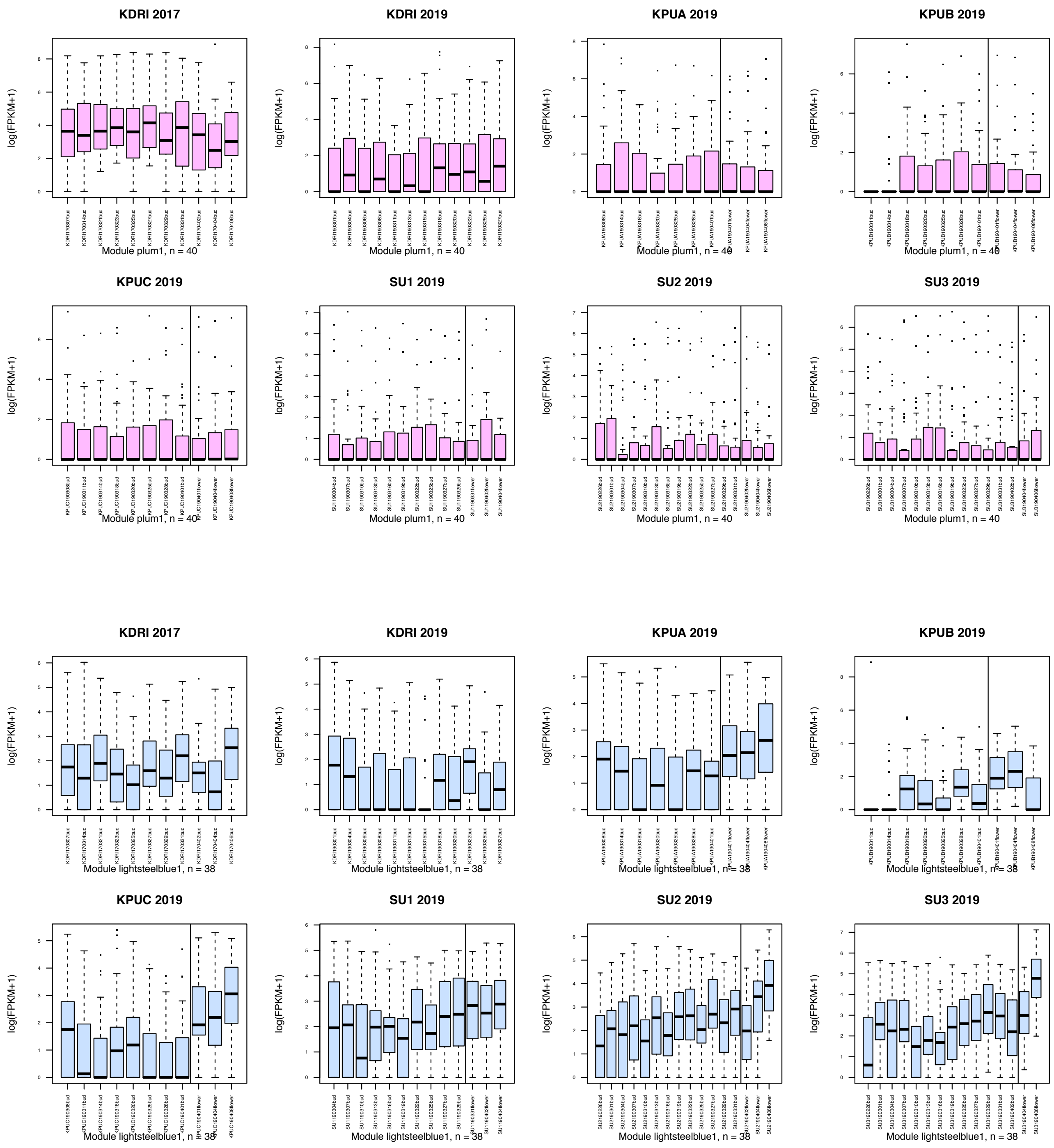


**Supplementary Figure S2** (Continued.)

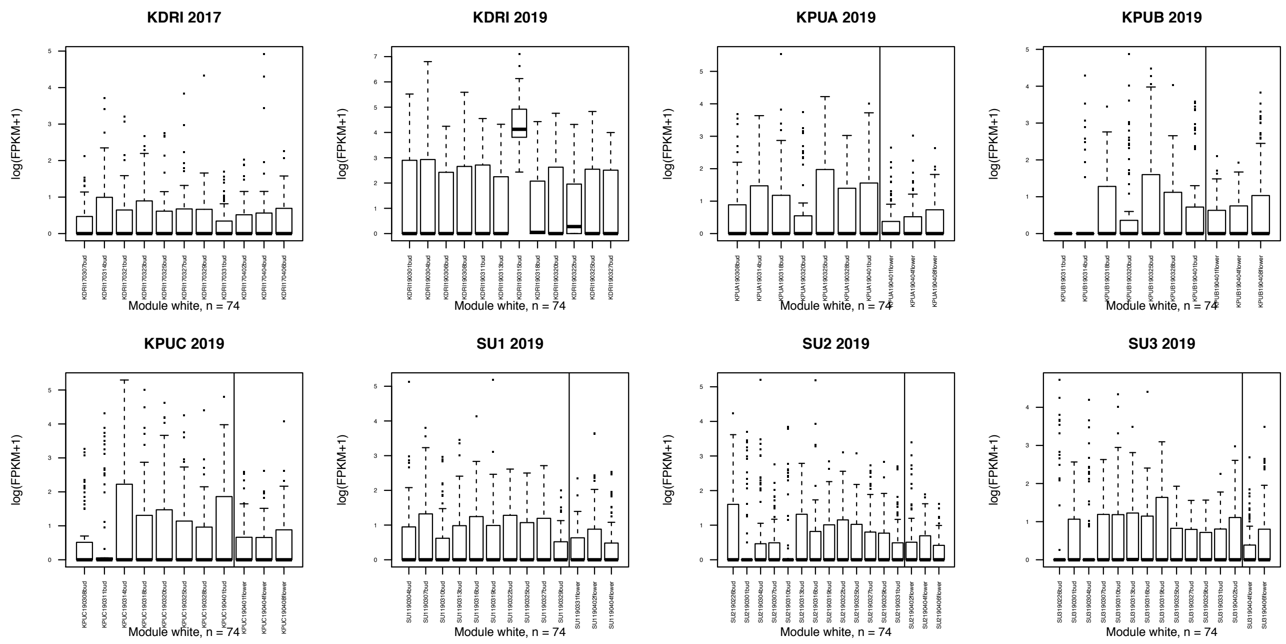


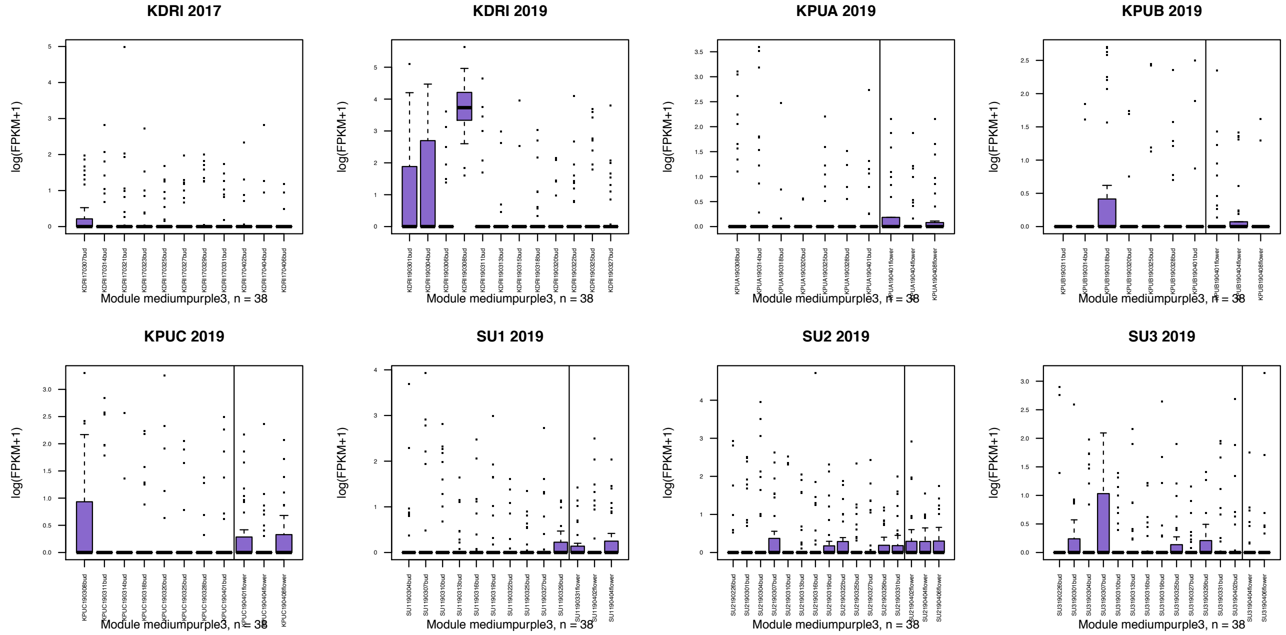


**Supplementary Figure S2** (Continued.)

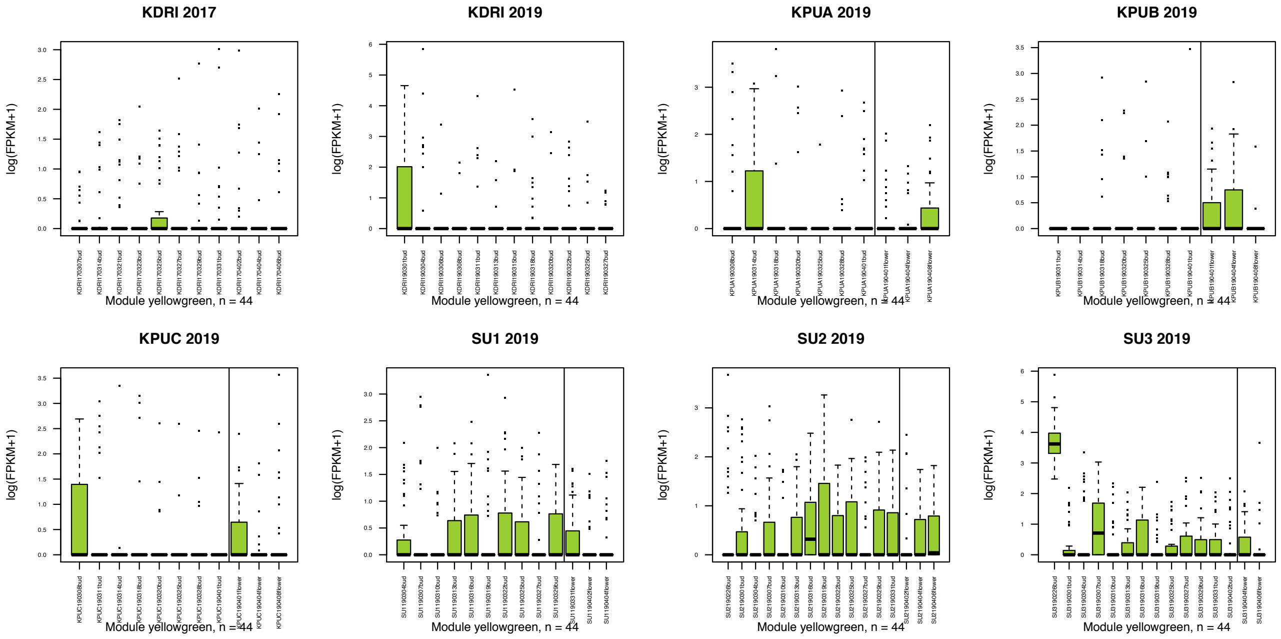


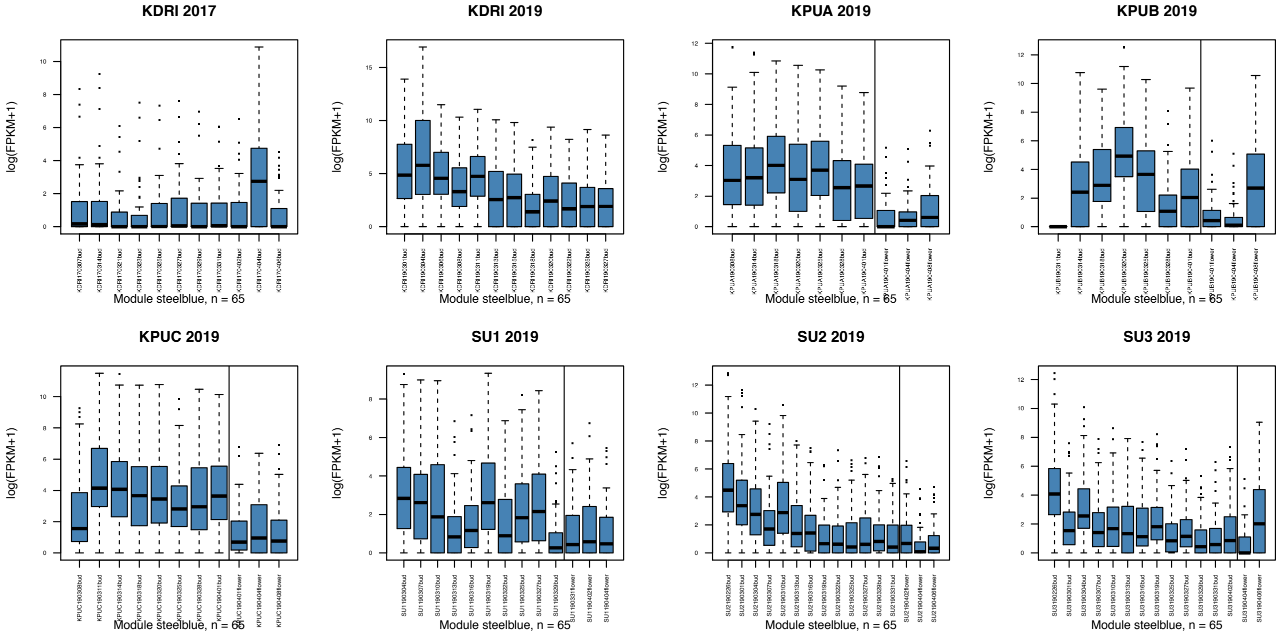


**Supplementary Figure S2** (Continued.)

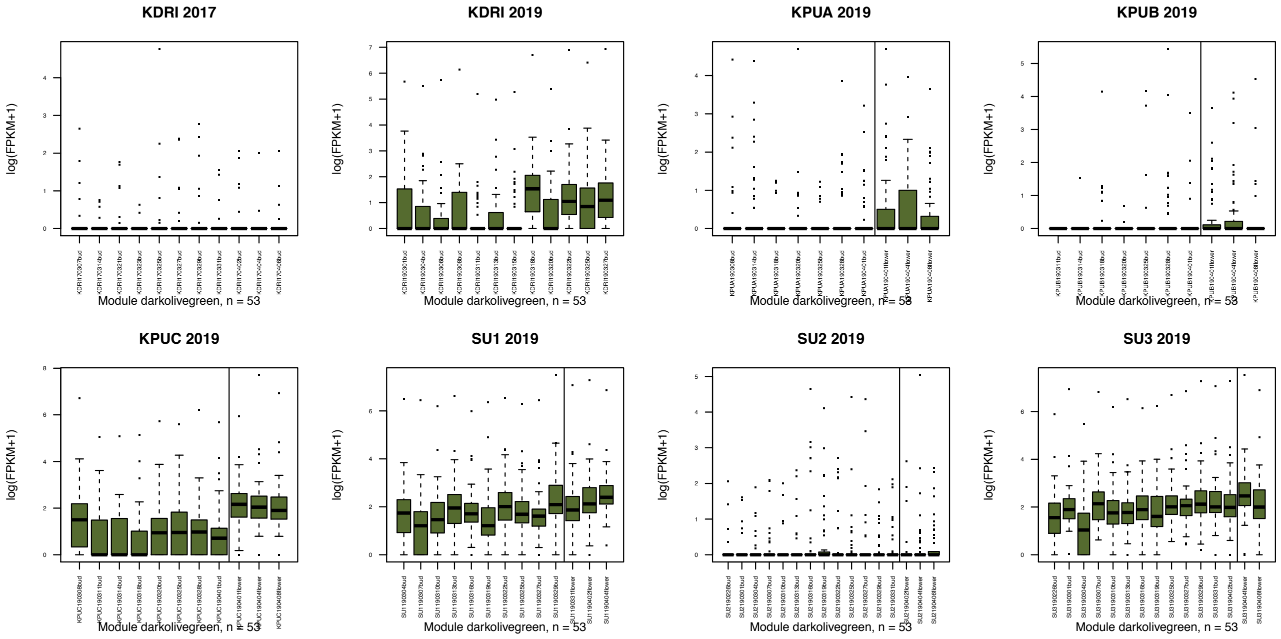


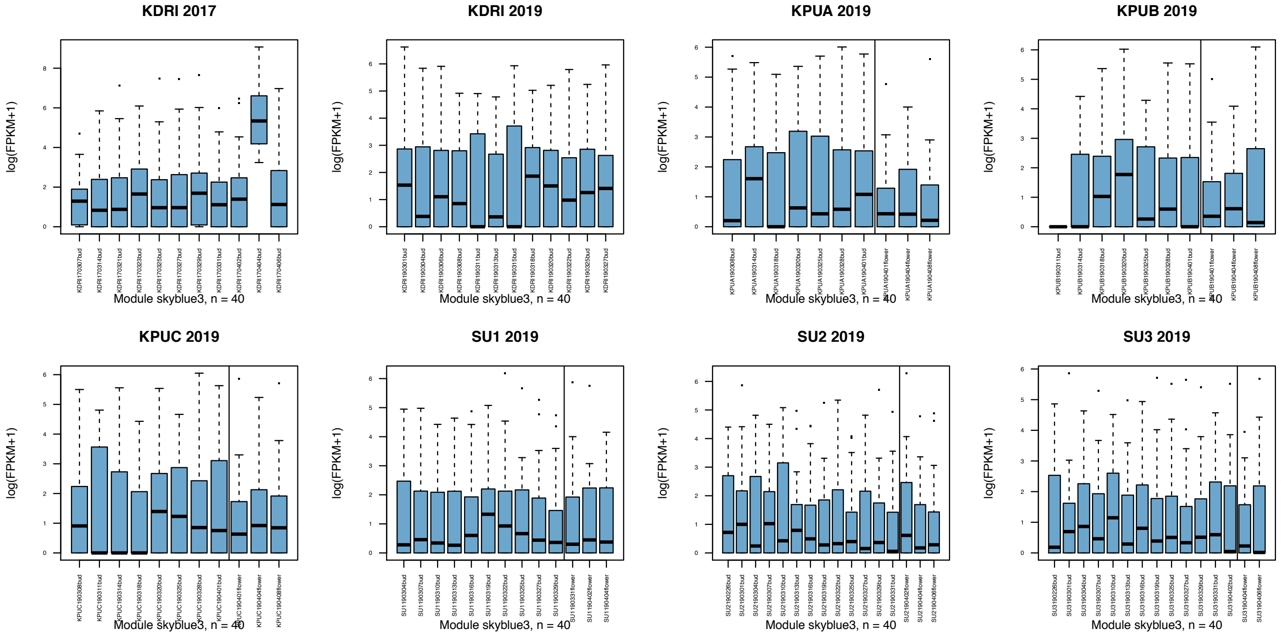


**Supplementary Figure S2** (Continued.)

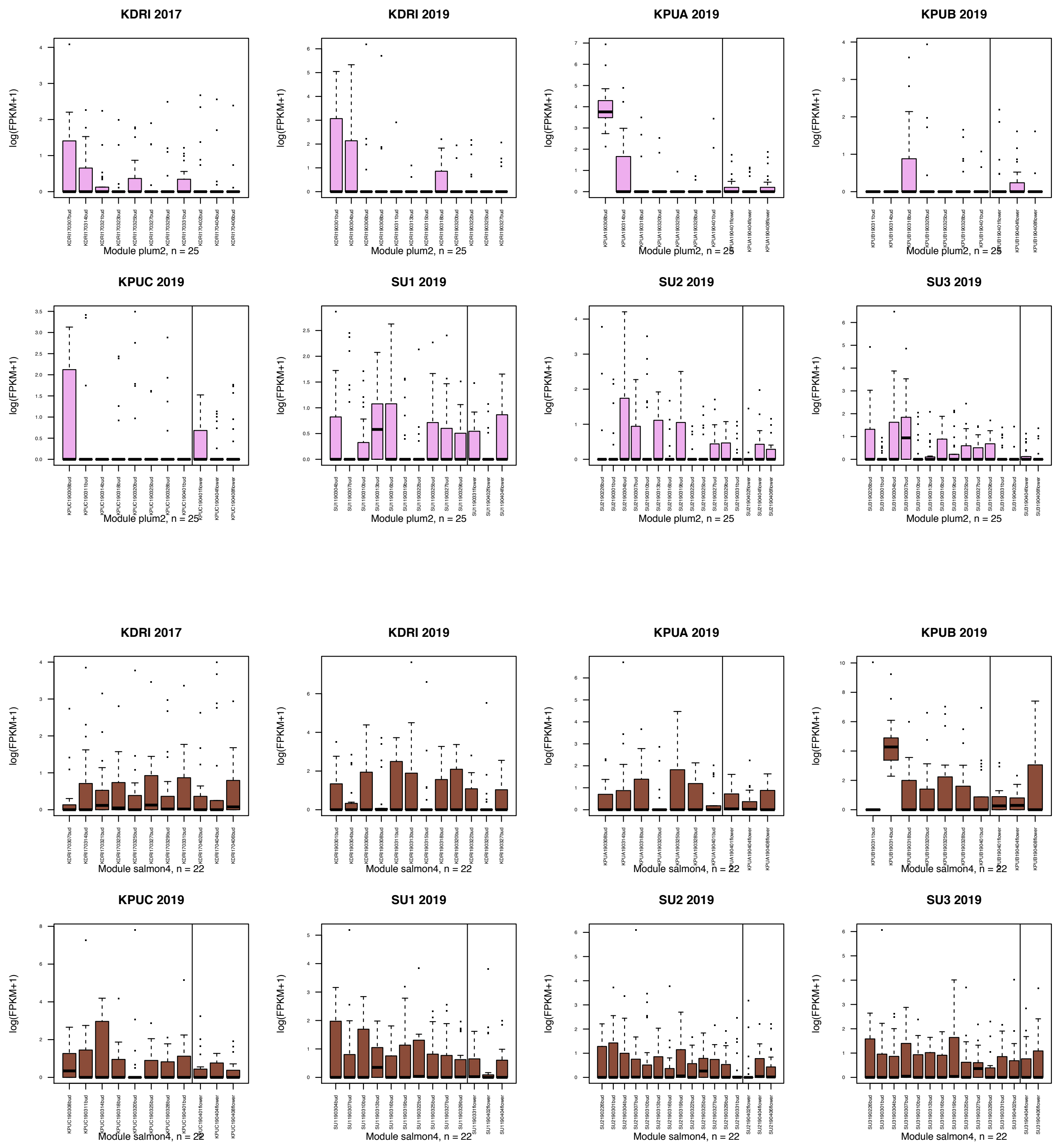


**Supplementary Figure S2** (Continued.)

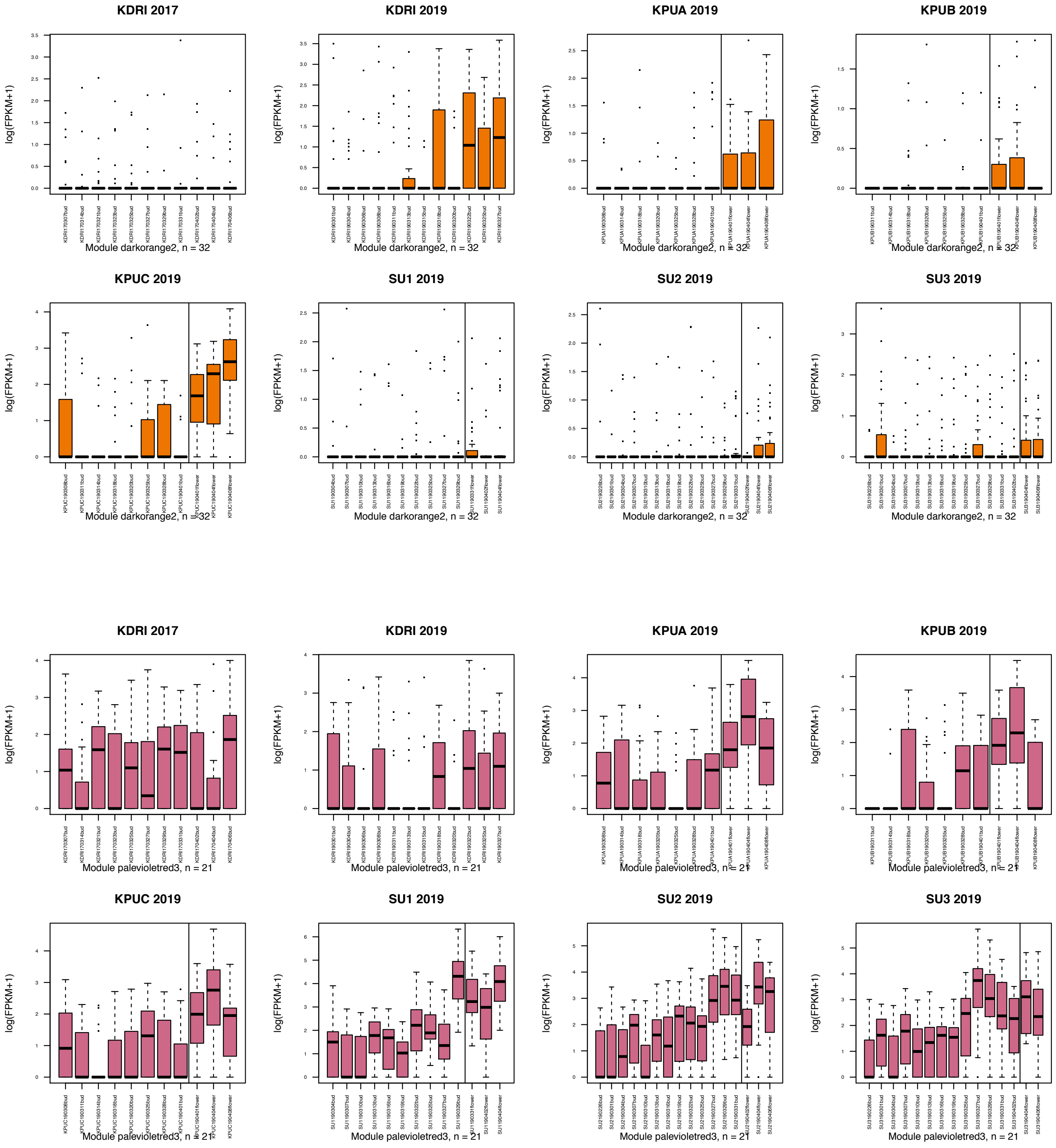


**Supplementary Figure S2** (Continued.)

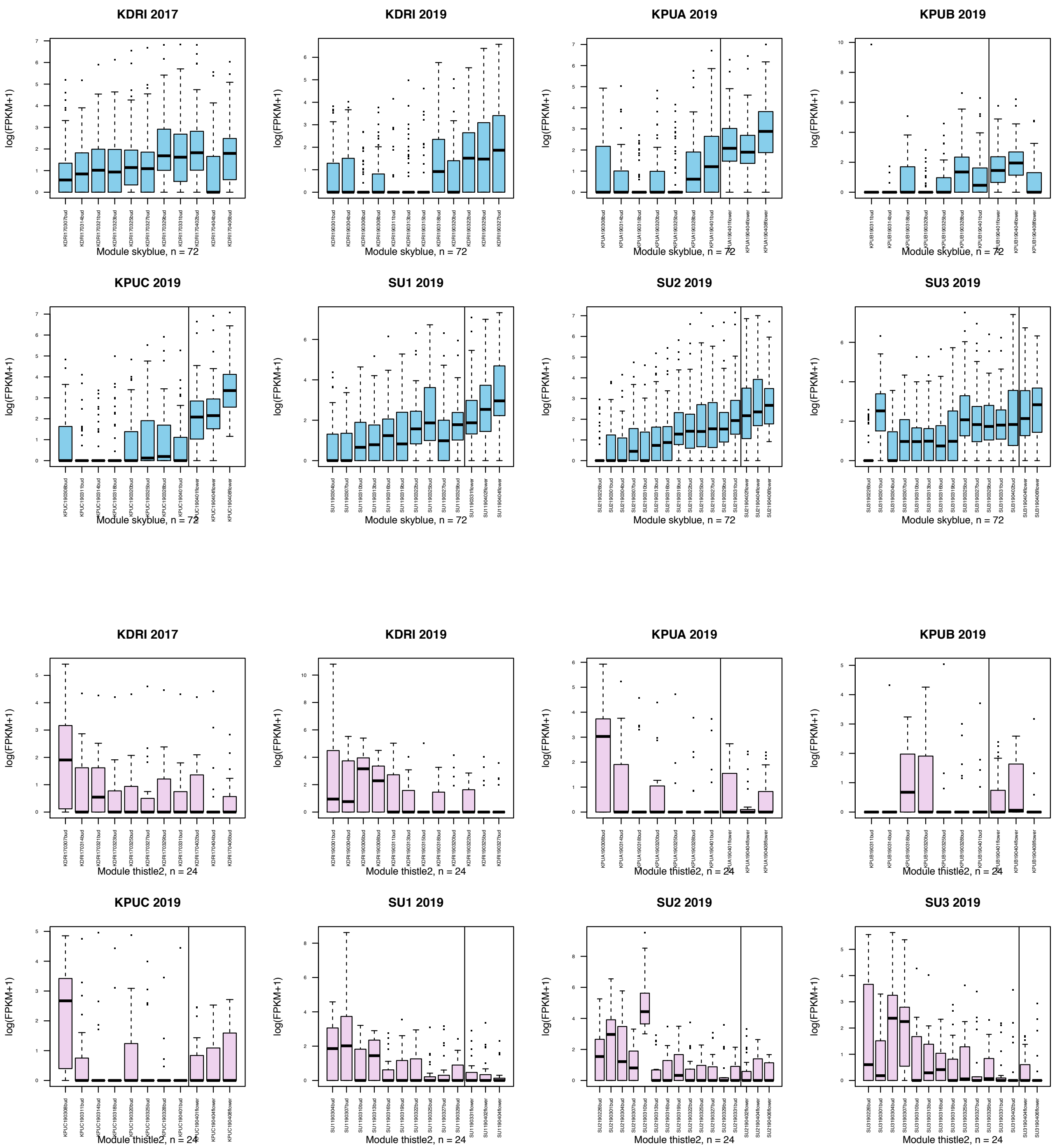


**Supplementary Figure S2** (Continued.)

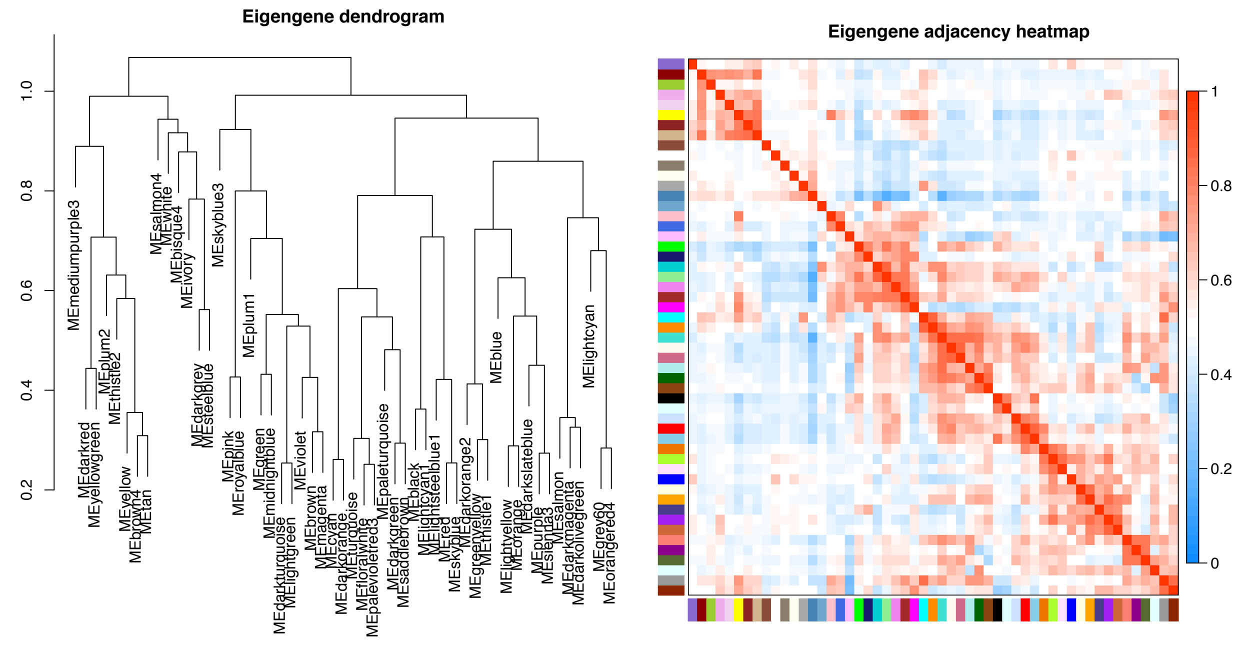


**Supplementary Figure S3** Dendrogram and heatmap of gene modules represented by eigengenes obtained by WGCNA.

#### GO terms enriched in the module "darkred" (4-5 WBF)

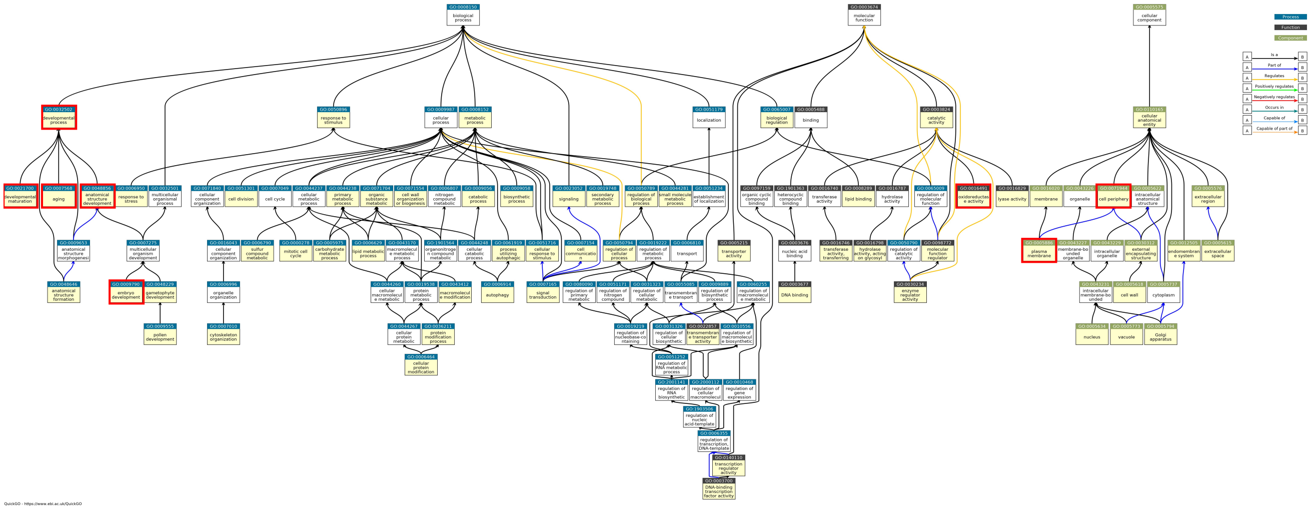

#### GO terms enriched in the module "tan " (4 WBF)

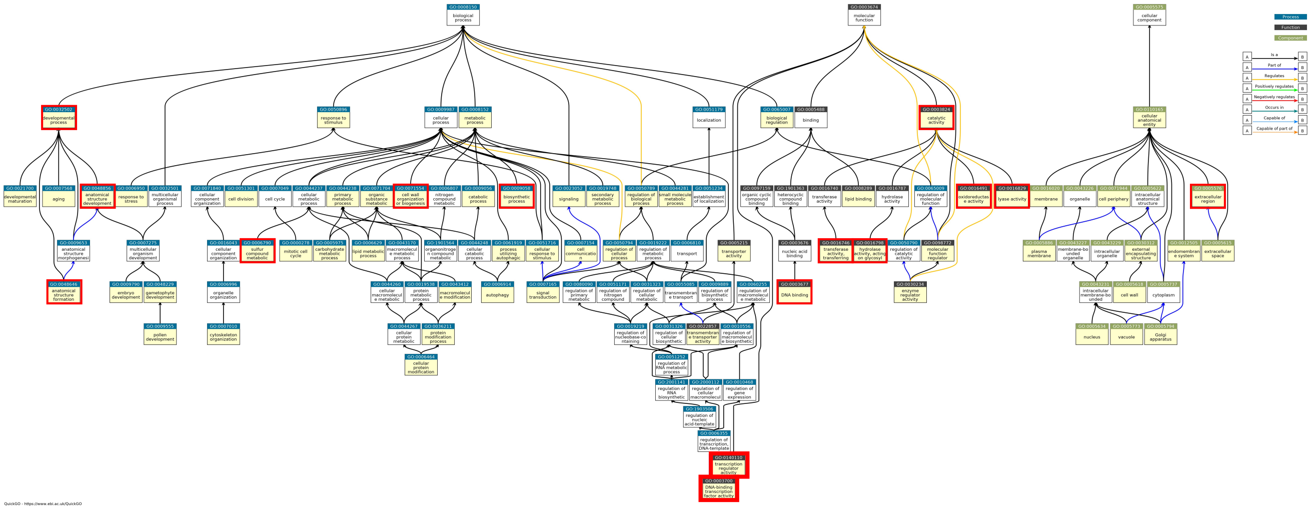

**Supplementary Figure S4** Gene Ontology (GO) hierarchy trees. Yellow boxes and red frames indicate GO terms enriched in all seven modules and in each individual module, respectively.

#### GO terms enriched in the module “pink” (3 WBF)

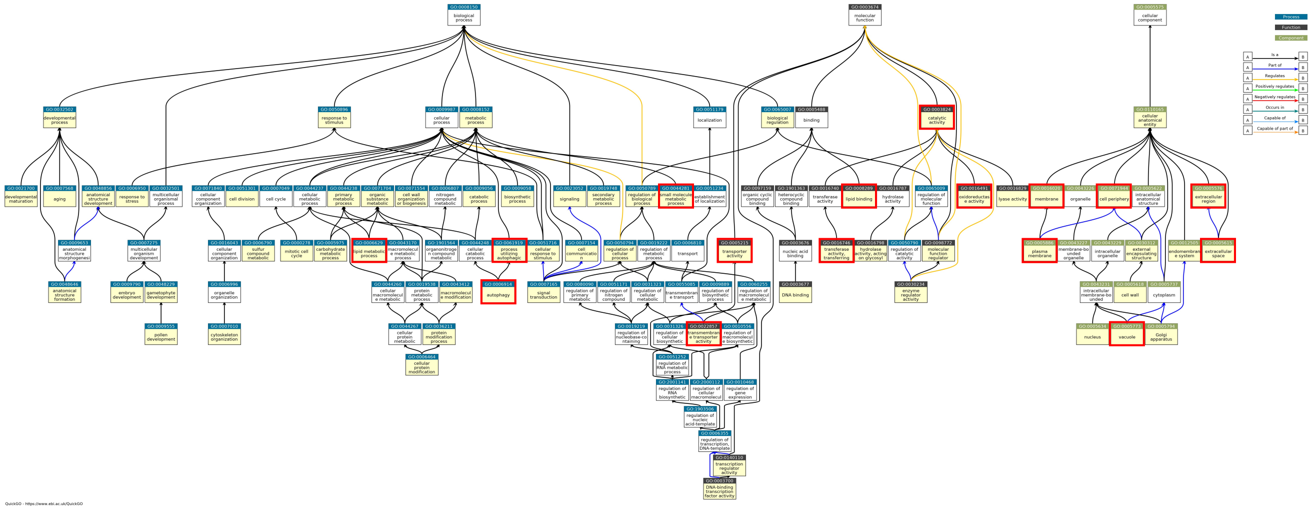

#### GO terms enriched in the module “royalblue” (2-3 WBF)

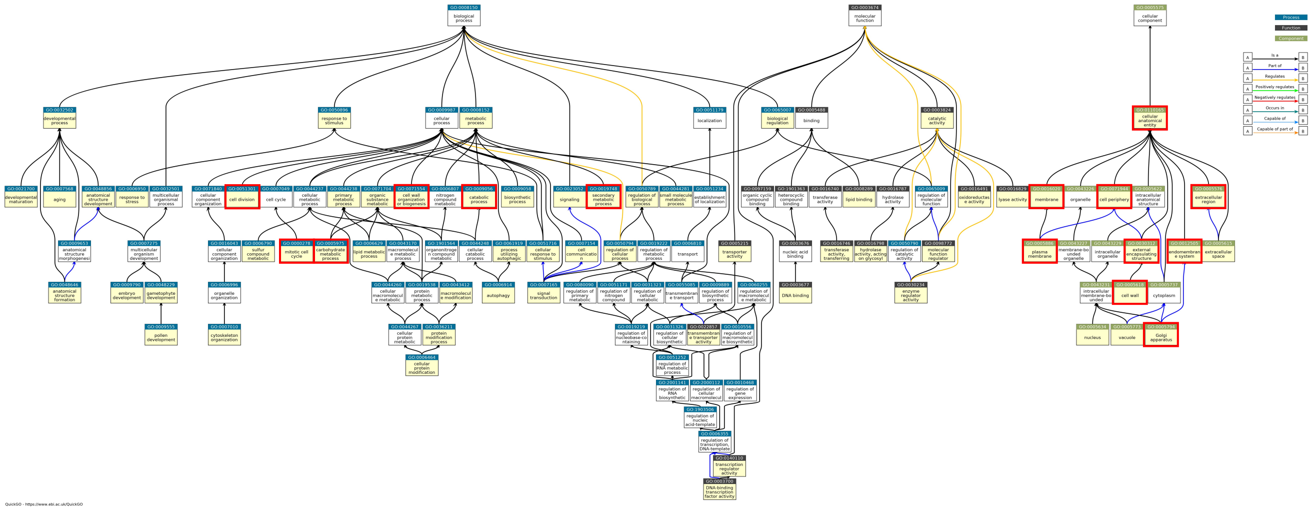

**Supplementary Figure S4** (Continued.)

GO terms enriched in the module “midnightblue” (1-2 WBF)

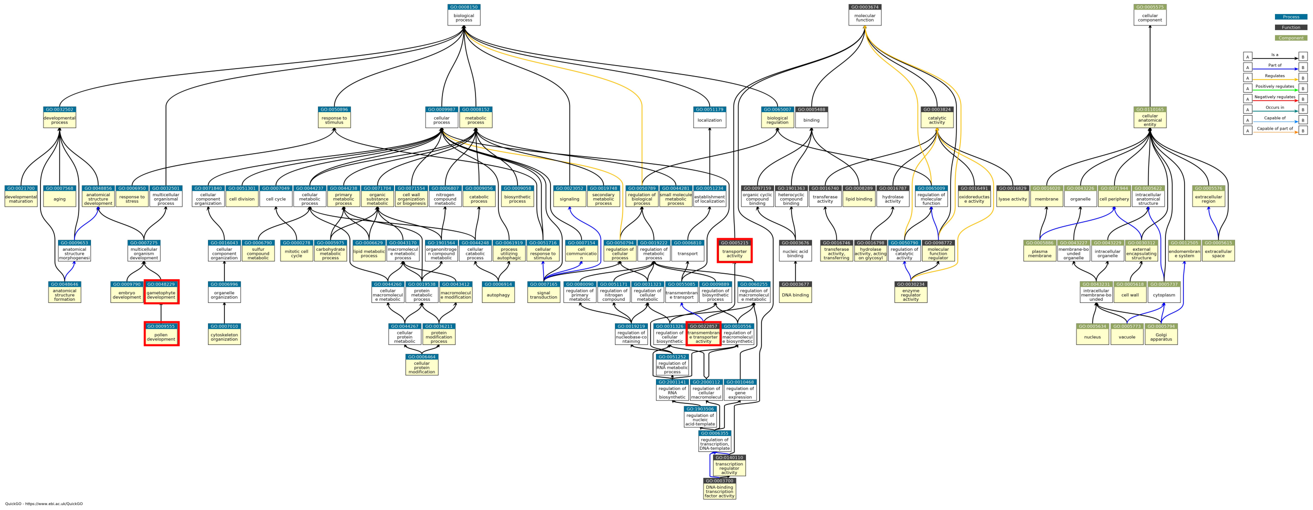

GO terms enriched in the module “black” (flowering day)

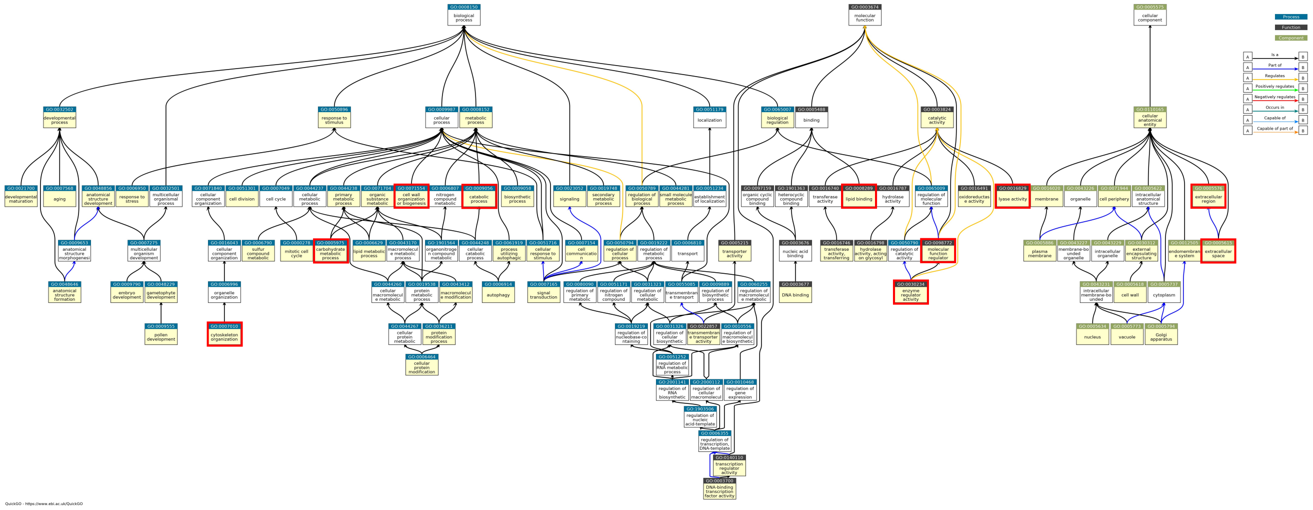

**Supplementary Figure S4** (Continued.)

### GO terms enriched in the module “skyblue” (1-2 WAF)

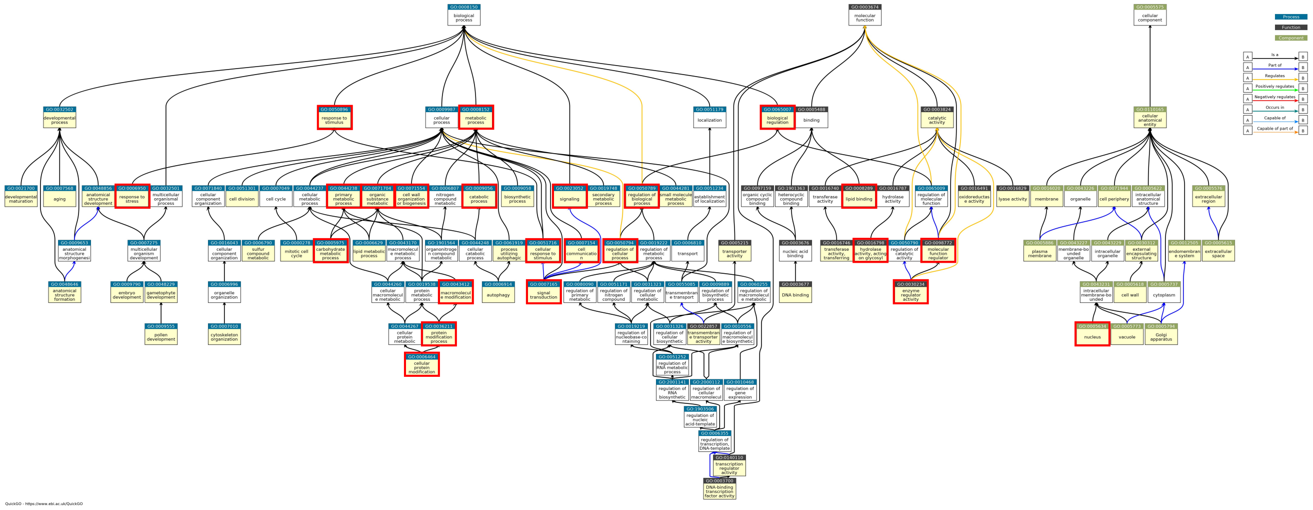

**Supplementary Figure S4** (Continued.)
